# Yeast growth responses to environmental perturbations are associated with rewiring of large epistatic networks

**DOI:** 10.1101/659730

**Authors:** Yanjun Zan, Örjan Carlborg

## Abstract

The phenotypic effects of genetic polymorphisms often depend on the genetic and environmental context. Here, we explore how polymorphic loci in large interaction networks contribute to complex trait variation and in particular how genetic effects and network topologies are influenced by environmental perturbations. This was done by reanalysing a dataset of >4,000 haploid yeast segregants grown on 20 different media. In total, 130 epistatic loci were associated with growth in at least one environment. The across-environment interaction network defined by these loci explained 69 - 100% of the narrow sense heritability in the individual environments. Environmental changes often resulted in network hubs being connected and disconnected from their interactors, leading to changes in additive effects of individual loci, epistatic effects of multi-locus interactions and the total level of genetic variance in growth. The largest variation in genetic effects across environments was found for epistatic loci that were highly connected network hubs in some environments but not in others. In environments where loci were highly connected, the segregating alleles epistatically suppressed or released genetic effects from their interactors. In environments where they were lowly connected, the same alleles made small or no contributions to growth. Hub-loci thus often serve as modulators, influencing the phenotypic effects of environmentally specific sets of interacting effector genes, rather than being effectors themselves. These findings illustrate the importance of the interplay between large genetic interactions networks and the living environment, both for individual phenotypes and population level metrics of genetic variation.

## Background

Most biological traits are influenced by the interplay of multiple genes and environmental factors. In most genetic studies, the phenotypic effects of segregating alleles are studied as static features. In reality they are, however, in many cases more likely to be dynamic due to influences by, for example, genotypes at other loci (G-by-G interactions or epistasis), variations in the environment (G-by-E interactions) or both (G-by-G-by-E interactions). Experiments have for many years found epistasis and G-by-E interactions to be abundant in many species^1–4^, and more recently it has also been possible to start exploring the environmental dependence of epistasis (G-by-G-by-E)^5–9^. For example, it has been shown how both pairwise ^10–14^ and high-order^15–17^ epistatic interactions can be conditional on environment. Consistent with this are results from studies of gene-interaction networks, where abundant rewiring has been detected in response to environmental changes^18–20^, although the phenotypic consequences of such changes are not yet known. Here, we evaluate the connection between the rewiring of gene-interaction networks, and changes in epistatic effects on complex traits, in response to environmental changes by reanalysing a large yeast dataset to answer the following two questions: *How often are high-order gene interaction networks rewired in response to environmental changes?* and *How does the environmentally induced rewiring of gene interaction networks influence the way that genetic polymorphisms contribute to complex traits?* By addressing these questions, novel insights could be gained on the connection between G-by-G and G-by-E interactions and how important this is for complex trait genetic variation.

Here, we reanalyse a public dataset of 4,390 yeast recombinant offspring (segregants) from a cross between a laboratory strain (BY) and a vineyard strain (RM). Every segregant was genotyped and phenotyped for growth in 20 environments (media)^21^. Previously, 939 additive QTL and 330 epistatic QTL were mapped when treating growth in these environments as independent traits^21^. In 11 of the 20 growth environments, large gene-interaction networks were identified where polymorphisms at highly connected epistatic network hubs supressed or released the genetic effects of their interactors^1^. Here, a complete interaction network was defined including all loci that contributed epistatically to growth in any of these environments. The rewiring of this epistatic network across the environments, and associated changes in how the interacting loci contributed to growth, were explored. Changes in connectivity of epistatic loci across environments often influenced their effects on growth. When changing the growth environment, network hubs were connected, and disconnected, from their interactions affecting their ability to epistatically suppress and release the genetic effects from radial loci. The interplay between epistasis and G-by-E interactions thus presents a mechanism by which populations can harbour hidden genetic variation in environmentally deactivated interaction networks for future selection response by network rewiring. The potential impact of these findings on the genetic maintenance of trait robustness in individuals and populations, the development of genetic models for quantitative traits as well their implications in areas where this might be of practical importance such as plant breeding and evolution, are discussed.

## Results

The yeast intercross population reanalysed here has earlier been analysed treating growth on different media as independent traits. Extensive epistatic interactions were then detected for growth on 19 of the 20 media^1,21^. Most epistatic QTL interacted with one or two loci, while a few interacted with several loci to form multi-locus radial networks^1^. Here, the data was reanalysed using a different approach (Figure 1). First, growth was considered a single complex trait with measurements in different media representing expression of the same phenotype in multiple environments. Differences in the content of the growth media constituted environmental perturbations. Second, the interaction networks mapped in each environment^1,21^ were combined into an across environment interaction network. A schematic example with four loci (A-D) in two environments (1-2) is illustrated in Figure 1A. Thereafter, the rewiring of this complete epistatic network across the environments, and the associated changes in the genetic effects, genetic variance and individual phenotypes were explored (Figure 1B). Here, we first present the complete interaction network and summarize its properties. Then, the rewiring of the interaction network across the environments is defined and how this connects to changes in the additive effects of individual loci, epistatic effects of multi-locus interactions and genetic variation of the population were explored. This was done by first using examples for subsets of environments and loci to illustrate principles that are later generalized in analyses of the complete network across all environments.

**Figure 1.**
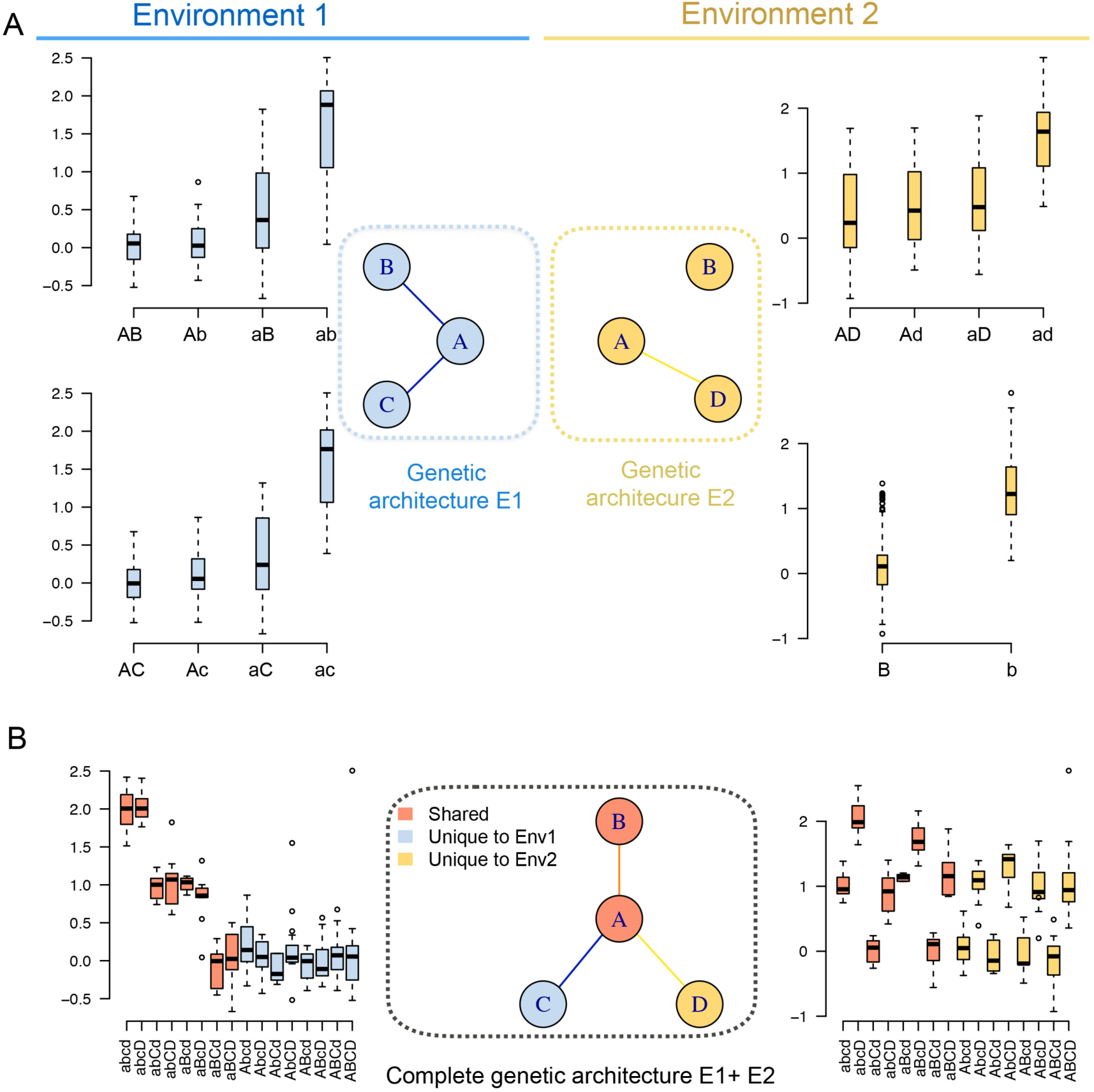
A schematic example of the analytical approach used to study the dynamic genetic architecture of haploid yeast growth in response to environmental perturbations. The circles illustrate loci (A-D) contributing to growth in a particular environment (blue/yellow) or in both environments (red). Lines connecting the circles indicate epistatic interactions between them. The box-plots illustrate the growth of the individuals relative to that in a standard environment (y-axis) with a particular single- or multi-locus genotype (x-axis). **A)** How loci contribute to growth in individual environments has been described in earlier studies of this population^1^. **B)** The known loci and interactions were used to define a complete across environment growth interaction network. The dynamics in the genetic regulation of yeast growth were studied by identifying the rewiring in this network, and associated changes in the genetic effects of individual and combination of alleles, across the studied environments.

## Defining an across environment epistatic network of growth loci

The complete across environment growth interaction network included 130 loci that were part of at least one of the 19 epistatic networks detected in the earlier analyses of this population^1^ (Figure 1A, Figure S1). Among these 130 loci, 38 (29%) had significant epistatic effects (Figure 2B) and 68 (52%) had significant additive effects (Figure 2C) on growth in multiple environments. The 130 loci were involved in 212 pairwise interactions, with the majority (88%) being unique to one environment, and 12% (27) being active in at least 2 environments (Figure 2D). Together, the 130 loci explained between 69-100% of the additive genetic variance in growth in the 20 environments (Figure 2E).

**Figure 2.**
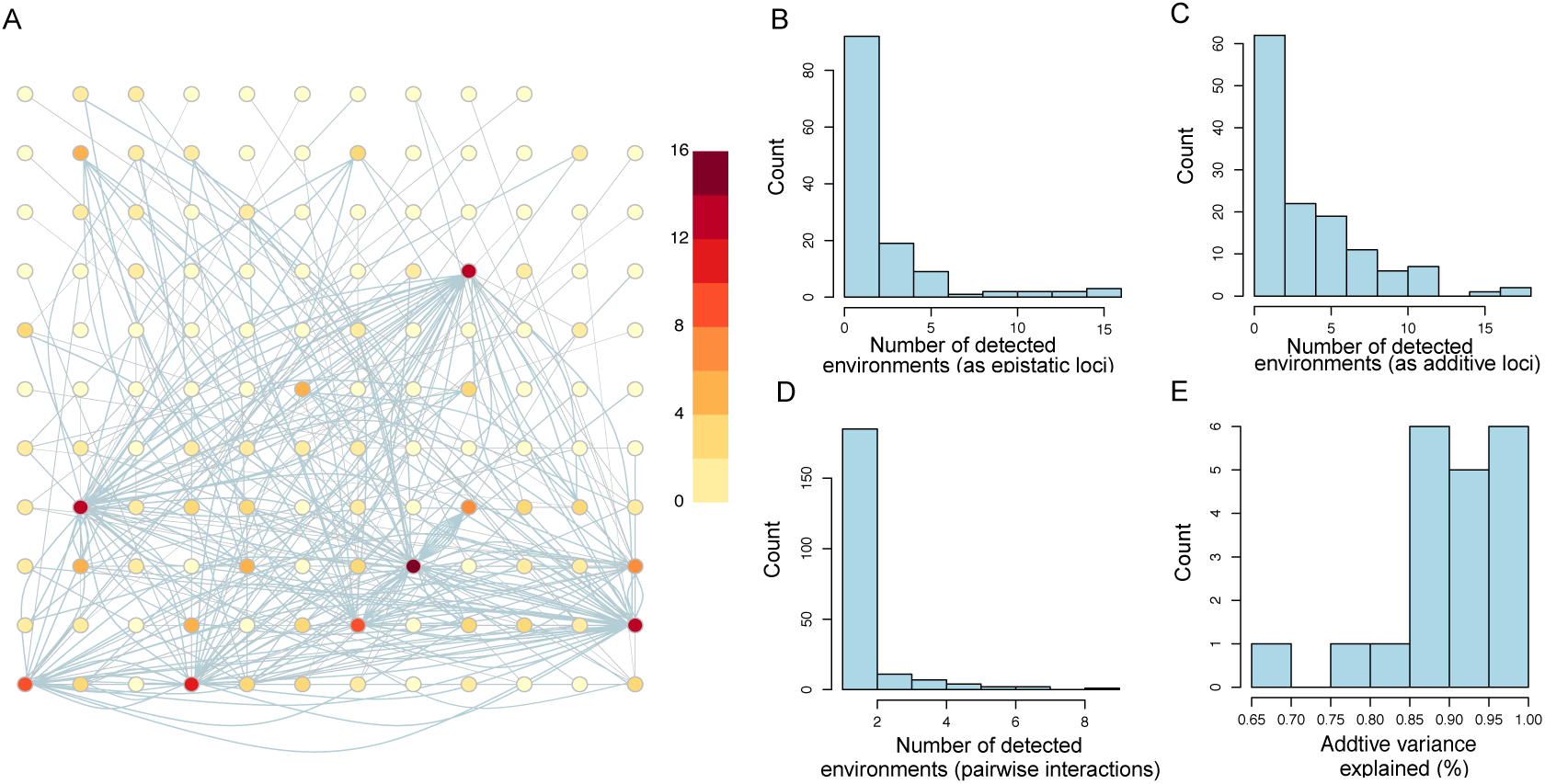
Summary of the features of the complete interaction network for yeast growth across the 20 studied environments. **A)** The complete across-environment interaction network includes 130 loci contributing epistatically to growth in at least one environment. Each node represents a locus, and its colour shows in how many environments it was involved in significant epistatic interactions. The edges represent significant pairwise interactions between loci, with the number of edges connecting pairs of loci corresponding to the number of times this pair was detected across the 20 environments. **B/C)** Histograms showing the number of environments in which the 130 loci were involved in at least one significant epistatic interaction/ additive effect. **D)** Histogram showing the number environments in which each of the 212 pairwise interactions were significant. **E)** Histogram showing how much of the additive variance for growth in the 20 environments that was explained by the 130 loci in the network.

The marginal effects of the loci in the network were thus more stable across environments than the interactions and a limited number of loci were highly interactive across many environments. In the following sections, we will explore the consequences of the extensive environmentally dependent rewiring of the epistatic network on growth. First, we use an example of a sub-network involving loci affecting growth across three environments to illustrate the principles and results, before later generalising them in analyses across larger networks and all environments.

## Associations between environmental perturbations, dynamics in the genotype-to-phenotype maps and epistatic network topologies: an example across three environments

The association between changes in the interaction network topology, and the changes of additive effects of individual loci, epistatic effects of multi-locus interactions and phenotypes resulting from particular combinations of alleles across environments, was explored for a 28-locus interaction network defined by connecting the epistatic loci detected for growth in 3 environments. These 3 environments were media where either Indoleacetic acid (IAA) or Formamide had been added, or where the carbon source was changed from Glucose to Raffinose (Figure 3A). Segregant growths were, overall, more similar on the media containing IAA and Formamide than in the medium with Raffinose as carbon source (PCA in Figure 3A; pairwise correlations Figure S2). In total, 6 (4) loci were detected with significant epistatic interactions in 3 (2) environments (Figure 3B). Consistent with the observations for the global network (Figure 2), the number of loci with significant additive effects was larger, with 9 (4) detected in 3 (2) environments (Figure 3C). Out of the 42 total pairwise interactions detected in the 3 environments, overlaps of 1 (9) pairwise interactions were observed for 3 (2) environments. The rewiring of the network associated with changes in the environment thus involved (Figure 3E-G) i) loci that were epistatic in all environments but with changes in the set of loci they interacted with, ii) loci that were epistatic in one environment and disconnected in the other, but that despite this contributed additively to growth or iii) loci that were epistatic in one environment and disconnected in the other, and that due to this no longer contributed to growth.

**Figure 3.**
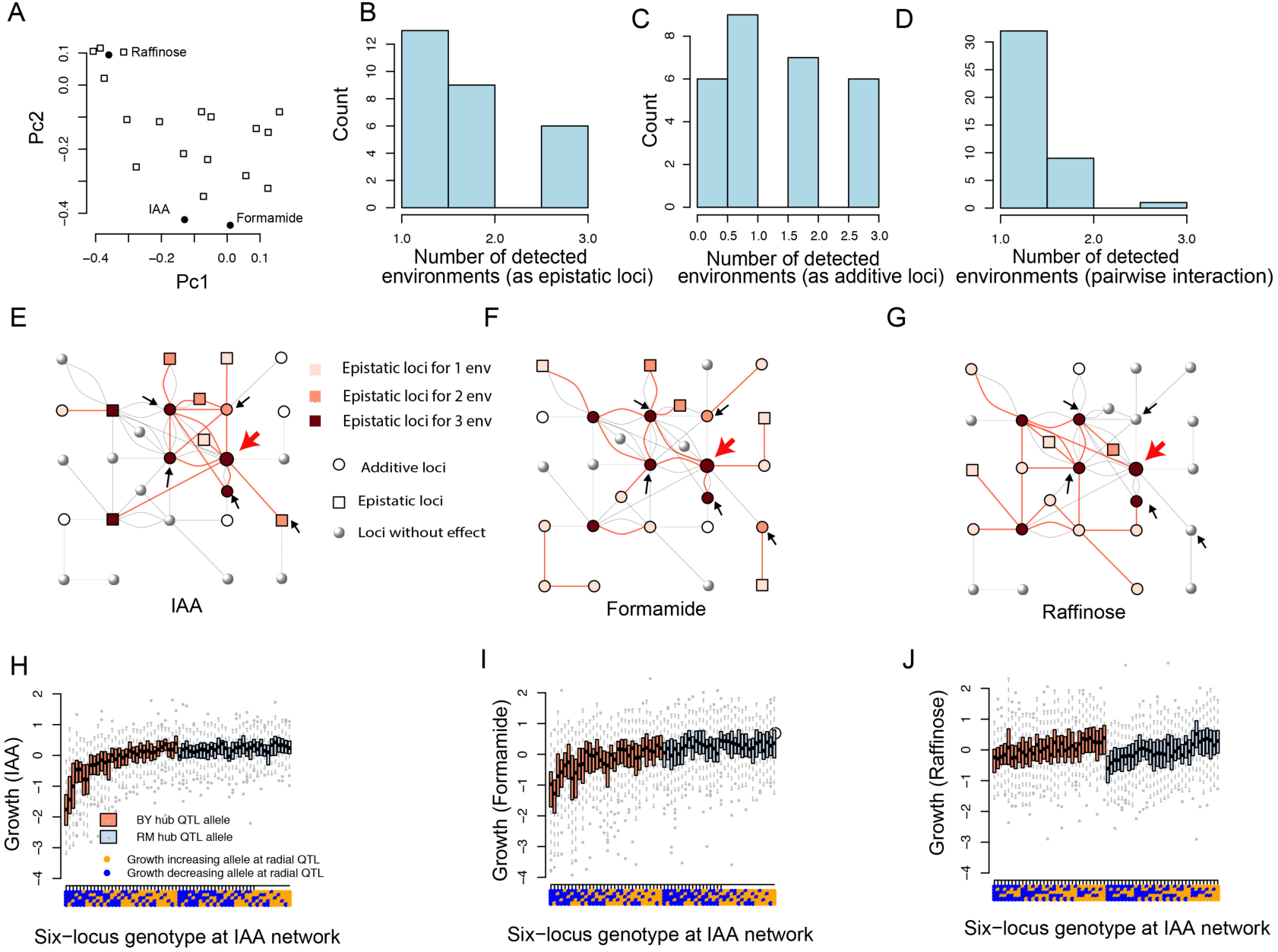
Illustrations of the relationships between environmental perturbations and contributions by high-order interactions contribute to yeast segregant growth. **A)** A two-dimensional PCA plot illustrating the resemblance in growths of yeast segregants across 20 media. Open squares represent a medium and the filled dots media with added IAA or Formamide or the medium where Raffinose was the carbon source. **B/C)** Histograms showing the number of environments in which the 28 loci had at least one significant epistatic interaction/additive effect. **D)** Histogram showing the number environments in which each of the 42 pairwise interactions were significant. **E-G)** Illustrations of differences and similarities in types of genetic effects (epistatic, additive or none) and network connectivity in the epistatic growth network on IAA, Formamide and Raffinose, containing media, respectively. The hub and radial loci in the IAA epistatic network studied earlier^1^ are highlighted with red/black arrows. The red lines connects significant pairwise interactions reported in Bloom et al^21^.. The grey lines indicate epistatic interaction detected in other media. Panes **H-J)** illustrate the genotype-to-phenotype maps for the six most significant loci in the highly connected epistatic network influencing growth on IAA (more details in Forsberg et al^1^) on IAA **(H)**, Formamide **(I)** and Raffinose **(J)**, respectively. They are illustrated with boxplots for each the 6-locus genotype. The color of the box indicates the genotype at the hub QTL (chrVIII: 98,622 bp; green/gray correspond to BY/ RM alleles). The x-axis gives the six-locus genotype class, where blue/orange dots indicate growth-decreasing/increasing alleles at the five radial QTLs (from top to bottom: chrXIV: 469,224 bp, chrIII: 198,615 bp, chrIV: 998,628 bp, chrXIII: 410,320 bp, chrXII: 645,539 bp).

Environmental changes were thus associated with extensive rewiring of the interaction network. To evaluate the connection between the rewiring and the effects of the loci on growth, a sub-network defined by the locus that was most extensively rewired was explored in more detail (Figure 3E-G). In total, 13 loci in the complete network contributed epistatically to growth in the medium containing IAA. One of these loci interacted with 7 other loci, defining an eight-locus radial growth network. The multi-locus effects of the six strongest loci in this radial network on growth in IAA-containing medium were explored in detail in earlier analyses of this data^1^ (Figure 3E). Here, we found a significant interaction between the 64 multi-locus genotype classes defined by these loci and the three environments (P < 2.2×10^−16^), suggesting a connection between the wiring of the network and the effects of the loci on growth. The highly connected hub-locus in this network was earlier shown to capacitate the effects on growth from the radial interacting loci^1^ (Figure 3H). When the segregants were grown on a medium containing Formamide, the connectivity in this six-locus network changed. Only 3 of the radial loci remained connected to the central hub. Three loci were rewired to interact with other loci, 1 no longer had any effect on growth and a new locus was connected to the hub (Figure 3 E/F). Also the genotype-to-phenotype map for the 6-locus network changed when the segregants were grown in media with Formamide. The main difference was a weaker capacitation effect of the hub-locus (Figure 3 H/I). Even larger differences in wiring, and the genotype-to-phenotype map, were observed when growing the segregants with Raffinose as carbon source. The hub-locus still contributed additively to growth, and was involved in one new epistatic interaction. It had, however, lost the connections to all the interactors contributing to growth on IAA, and two of these no longer contributed to growth. The disconnection of the hub-locus from its original interactors likely explains the loss of phenotypic effect on growth in this medium (Figure 3J). The gradual deactivation of the capacitation (epistatic) effect of the hub-locus from IAA->Formamide->Raffinose was also consistent with the phenotypic correlations of segregant growths on these media as illustrated in the PCA plot (Figure 3A) and the pairwise correlations of growths in the media (Figure S2). Together these results illustrate the association between the rewiring of the epistatic network and changes in growth effects of the loci involved across environments, i.e. how the connectivity of the hub-locus defines its ability to act as a capacitator of other growth loci.

### Extensive network rewiring for loci with variable contributions to growth across environments: effects across all environments

To generalize the findings from the three-environment example above, the connectivity and contribution by all networks to growth across all 20 environments were analysed. The results are described in detail in the sections below.

#### Rewiring of high-order gene interaction networks in response to environmental changes is common for highly connected loci

In total, 13 loci with more than 4 epistatic interactors were detected in at least one of the 20 environments. The networks defined by these 13 hub loci and their interactors included 70% (91 of 130) of the loci in the complete network (Figure 2A; Figure S3). The connectivity of these 13 networks was highly dynamic across the 20 environments, with hubs being fully connected in some environments but completely disconnected in others (Figure S4A; Figure 4A). Although epistatic network connectivity changed across environments, it was common that some of the interactors (0-83%; median 30%) still had significant additive effects in other environments (Figure 4B; Figure S4 B). The detected epistatic interactions thus seemed to be more sensitive to environmental perturbations than additive effects (Figure 4 A/B; Figure S4 A/B). In addition, when the interactions between hub and radial loci were disconnected from one environment to another, the total contribution by the whole network to the growth variation decreased (Figure 4C). The rewiring of the epistatic networks in response to environment changes was thus associated with their contribution to growth.

**Figure 4.**
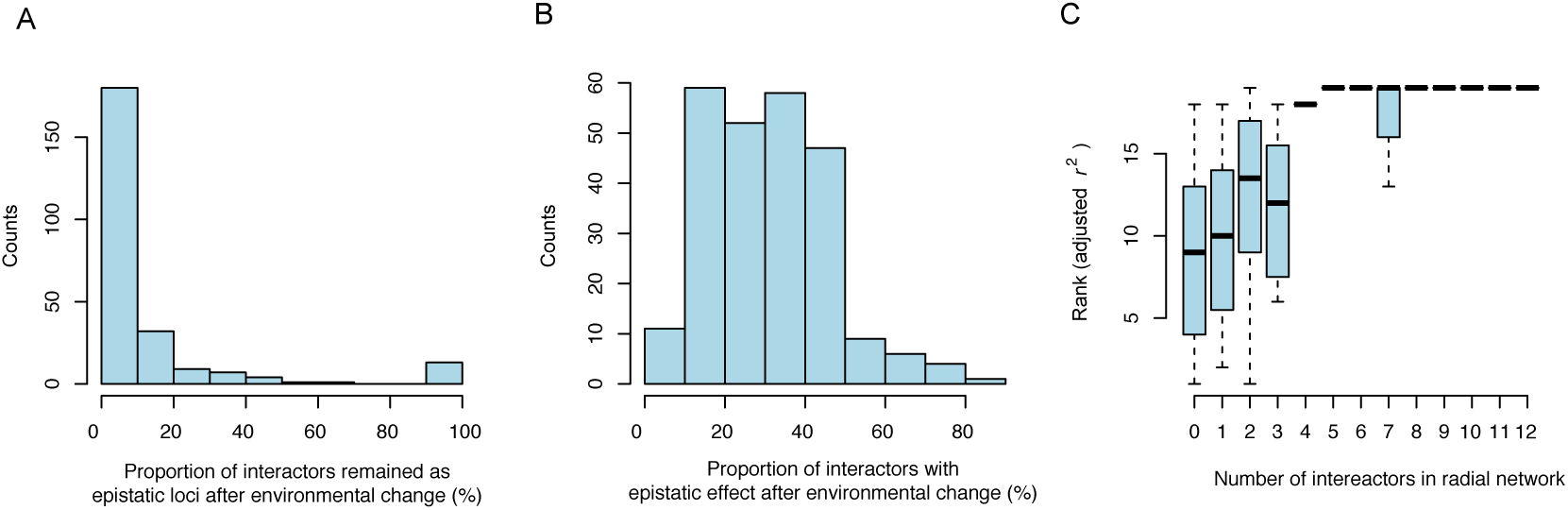
Illustrations of the dynamics of the largest mapped epistatic networks, and their contribution to growth variation, across all tested environments. **A)** Histogram illustrating the changes in connectivity of 13 epistatic networks, with more than 4 interactors, across the 20 environments. For each network, a set of interactors was defined to include the radial loci in the most highly connected environment. The percentage of these interactors that were epistatically connected to the hub in each of the remaining 19 environments was calculated (x-axis). The overlap of interactors in the 13 networks across the 20 environments was summarized as the counts of environments with similar percentages of shared interactors (y-axis). **B)** Histogram illustrating the percentage of interactors (defined as in A above) that have significant additive effects across the 13 networks and 20 environments. The x-axis shows the percentage of these interactors that have additive effects across the other 19 environments. The y-axis summarizes these percentages across the 13 networks. **C)** In each environment, adjusted r^2^ values are calculated for all networks and ranks of these model fits were assigned. The association between the connectivity of the epistatic networks (x-axis; number of loci connected to the hub), and their contributions to the variance in growth (y-axis; rank of adjusted r^2^ values) across the 13 networks and 20 environments, is illustrated as box-plots of these ranks grouped by the number of interactors.

### Variations in marginal additive effects across environments associated with rewiring of epistatic networks

Significant additive effects on growth in at least one environment were detected for 311 loci in the genome. Although not every locus had significant effects on growth in every environment, these loci did as a group contribute significantly to growth in most environments (Materials and Methods; Table S1). The effects were generally stronger in one or a few of the environments (Figure S5). For example, more than half of the loci were unique to one environment and only 9 loci were associated with growth in more than 10 environments. QTL by environment interactions were thus abundant with 98% (307) loci displaying statistically significant QTL by environment interaction after multiple testing corrections (Figure 5; Table S2).

**Figure 5.**
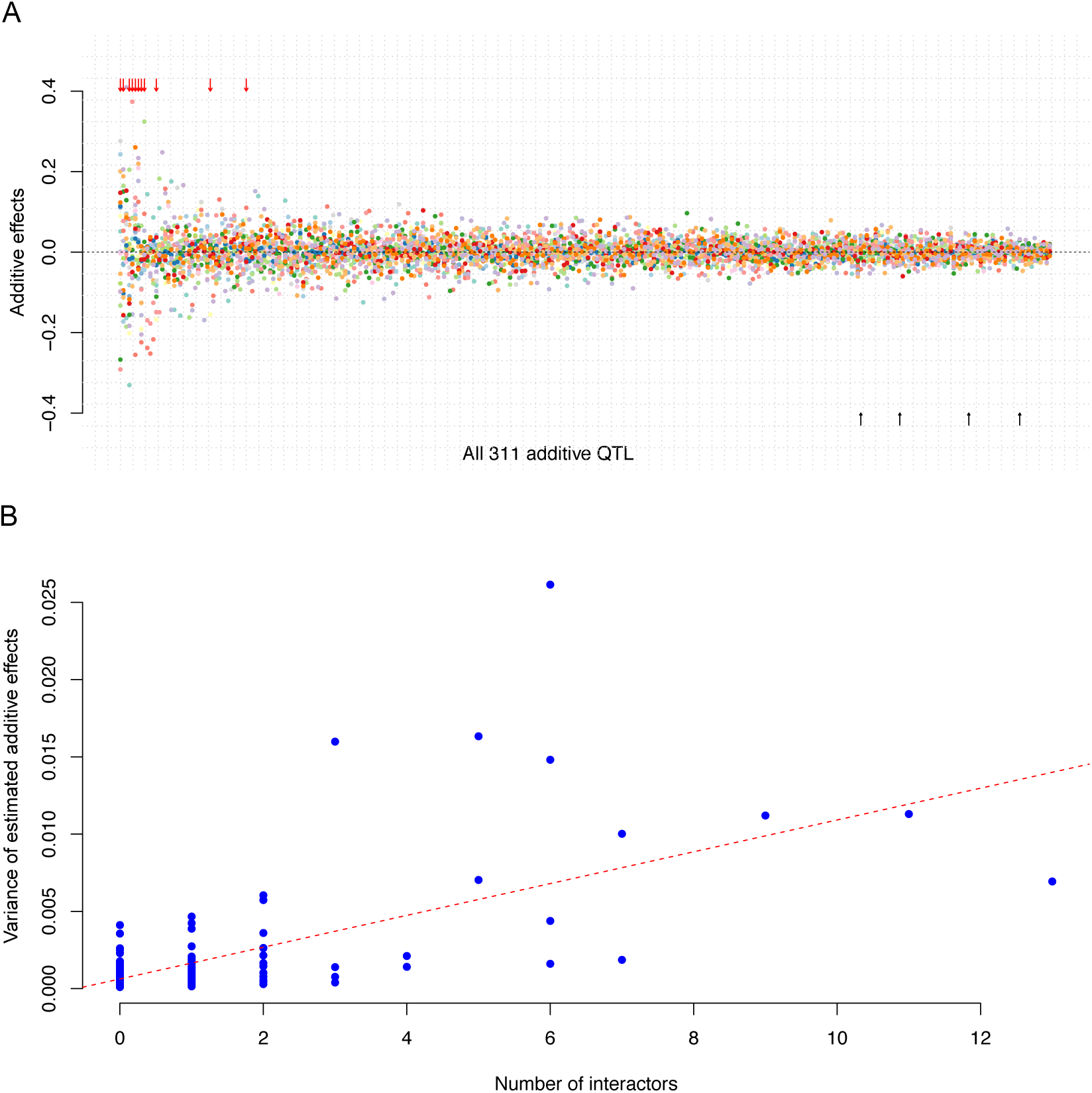
The variations in additive effects on segregant growth and their associations with network connectivity. **A)** 311 loci had significant additive genetic effects on growth in at least one of the 20 environments. The estimated additive effects (y-axis) for each locus (x-axis) in the 20 studied environments are illustrated using dots in different colours. The loci (represented by the most significant SNPs) are sorted from left to right by the variance of the estimated additive effects across the environments. All additive effects estimates were obtained by fitting all 311 loci jointly in a linear model. The 4 loci without significant genotype by environment interactions are indicated with black arrows below the x-axis. Out of the 13 highly connected loci (interacting with more than 4 other loci), 11 had significant additive effects in at least one environment^1^ and they are highlighted with red arrows on the top. **B)** An illustration of the relationship between the maximum number of epistatic interactions (x-axis) and the variance in their estimated additive effects across the 20 media. Each blue dot represents a locus with a significant additive effect. The red dashed line is the regression line (P value = 7.1 × 10^−39^).

All 11 QTL that were highly connected (> 4 epistatic interactors) in at least one environment displayed large variations in the marginal additive effects across the environments (Figure 5A; P-value = 6.1 × 10^−8^; Wilcoxon rank sum test). As the network wiring of these loci also changed much between environments, we hypothesised that the additive effects in the different environments were associated with changes in the number of G-by-G interactions of the locus. To test this, we estimated the correlation between the variations in the number of epistatic interactors and variance in the additive effects across the 20 environments. This correlation was highly significant (Figure 5B; P value = 7.1×10^−39^; linear regression), suggesting that the variation in additive effects of polymorphisms across the 20 environments was connected to the rewirings in the epistatic network.

### Variation in the non-additive genetic variance contributed by loci across 20 environments is correlated with the rewiring of the epistatic network

As reported earlier^21^, there were considerable variation in both the phenotypic and genetic variation in growth (Figure S6A), as well as broad sense (0.11 < H^2^ < 0.88; median 0.64) and narrow sense (0.09 < h^2^ < 0.70; median 0.43) heritability between environments (Figure S6B). The contributions from the epistatic network rewiring to the differences in the non-additive genetic variances between the environments were evaluated across all traits and loci. A significant positive correlation (P-value = 5.6 × 10^−31^; regression; Figure 6) was detected between the differences in the levels of non-additive genetic variance (H^2^_env1_-H^2^_env2_) and differences in the number of total epistatic connections. This highlights the general importance of G-by-G-by-E interactions for growth in this population, as well as the contributions by the high-order network connectivity for the non-additive variance in particular.

**Figure 6.**
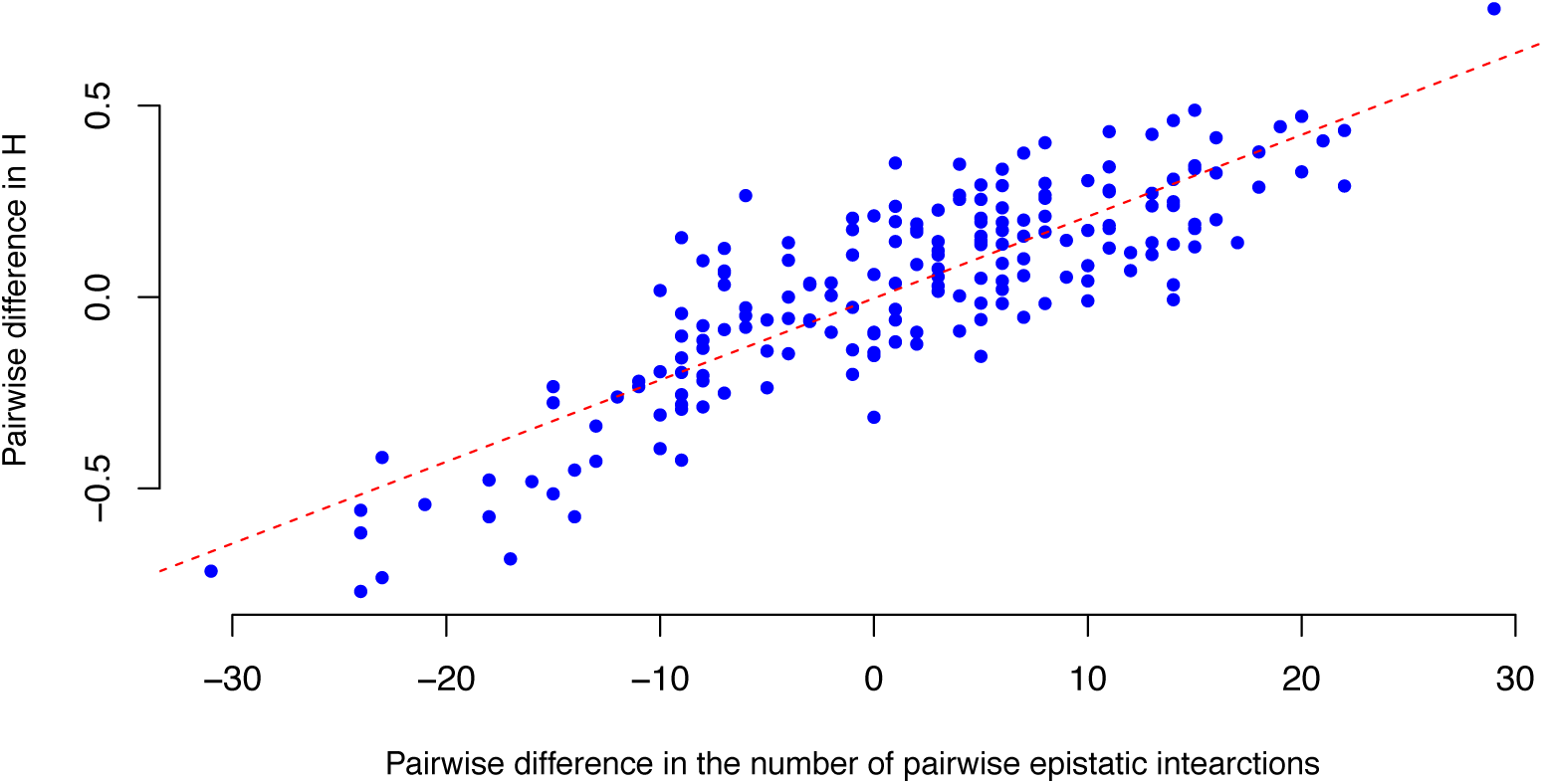
Illustration of the relationship between the network wiring and amount of non-additive genetic variance explained by the epistatic network across the growth environments. A significant correlation (P-value = 5.6 × 10^−31^ from regression) was detected between the pairwise differences in the number of epistatic interactions (x-axis) and the non-additive variances (H^2^_env1_-H^2^_env2_; y-axis) for the loci in the complete interaction network across the 20 environments.

### Discussion

Yeast has, for many years, facilitated studies on the genetic basis of complex traits. Large haploid segregant populations can be generated, where alleles segregate at intermediate allele frequencies. Earlier work using such populations have shown their value in, for example, dissecting the genetic basis of gene expression^22^ and understanding the sources of the “missing heritability”^23^. By generating very large populations, and high-throughput phenotyping the segregants in replicates across many environments, such populations have also facilitated studies on more complex genetic mechanisms including genotype by environment interactions^1–4^, how high-order interactions contribute to complex trait variation and prediction^21^ and the conditional effects of epistatic interactions on environmental factors^7,9,19^. Here, we reanalysed a very large, publically available haploid segregant yeast dataset where one complex trait – colony size – had been measured across many environments. In this way, we could reveal that the gene-interaction networks contributing to growth, and their associated high dimensional genotype to phenotype maps, were highly dynamic across growth environments. Further, we also illustrated the importance of these dynamics for the differences in trait expression of individual segregants, and genetic variance expressed in the population, across the evaluated environments.

### The effects of genotype-by-environment interactions on yeast growth

The studied dataset did not allow direct estimation of the contribution by systematic environmental (i.e. growth medium) effects on the phenotypes as the available growth measurements were pre-normalised against growth in a control medium^21^. However, estimates of the contributions by direct genetic (G) and genotype-by-environment (G-by-E) effects showed that direct genetic effects only contributed about 1/3 as much to the variance in growth compared to contributions by G-by-E. Consistent with the large contributions from G-by-E, we found large differences in the genetic architecture of growth (defined as associated loci and their genetic effects) across environments. The wiring into interaction networks, as well as marginal additive effects, of many growth loci were highly environmentally specific. Likely as a consequence of this, the phenotype changed across environments with the rewiring of the genetic growth network. The same polymorphisms thus contributed to growth in different ways across the environments, regardless of whether their effects were quantified via marginal additive effects or variance explained by the complete network. In the following sections we discuss these findings in more detail as well as their implications for the mapping, and understanding the evolution of complex traits.

### High environmental specificity for both additive and epistatic genetic effects

Most of the detected loci, regardless of whether they were originally mapped via their additive effects or epistatic interactions, only contributed to growth in one or a few environments. This in itself indicates extensive G-by-E interactions for growth in this population. However, loci mapped by their additive effects replicated to a greater extent across environments than loci mapped via their epistatic effects. Epistatic loci were thus, often highly environmentally specific suggesting a connection between G-by-G and G-by-E in this population. This is consistent with previous analysis on a similar yeast cross, where cryptic genetic variation is released under rare allelic combinations in specific environmental conditions^9^. Here, this G-by-G-by-E was quantified by evaluating the rewiring of the interaction network across environments, and its importance for growth was studied by measuring the associated changes in the allelic effects on growth. The low overlap of epistatic interactions across environments (Figure 2, Figure S1) illustrates the dynamics in the interaction network in response to environmental perturbations. The epistatic network is almost always rewired between the environments, likely in response to the different stressors in the media. This illustrates the role of G-by-G-by-E interactions for yeast growth where the environmentally induced rewiring influences the genetic effects of individual and combinations of loci, altering the level of genetic variance of the population in the environments. Such dynamic variations in the allelic effects on growth across environments is thus likely important in many studies of complex traits, for example when aiming to understand the processes allowing individuals and populations respond to changes in the surrounding environments.

### Environmental specificity of high-order interaction effects facilitates buffering of genetic effects in populations

Capacitating epistasis, defined as interactions where major loci in epistatic networks that can hide/release the genetic variance of other interactors, in radial multi-locus networks has earlier been identified as an important mechanism by which polymorphisms in one or a few hub-loci can buffer a significant amount of genetic variation from its interacting loci^1,3^. Here, it was shown that the topology of the interaction networks, as well as the capacitation effects of the hub-loci, generally were environment specific: the capacitors released the large phenotypic effects of its interactors in some environments, whereas they were often disconnected from them in others. Together with previous findings where yeast growth plasticity was found to be regulated by environment-specific multi-QTL interaction^7^, G-by-G-by-E interactions are thus likely to be a buffering mechanism allowing populations to accumulate cryptic genetic variation in a wide range of environments, for later release in response to environmental perturbations facilitating large and rapid responses.

### Network capacitation influences individual robustness to environmental perturbations

Our results suggest that the interplay between network interactions and environmental factors are important also for individual robustness. Individuals with more non-capacitation hub alleles perform better, on an average (Figure S7), and tend to show less variability across environments. This might be of relevance to, for example, plant breeding where one of the key challenges is to minimise the impact of genotype by environment interactions on production. Interactions in large gene interaction networks, buffered by environmental factors, might thus be an important driver of observed G-by-E interactions in populations. Targeted breeding for particular alleles at central network hubs might provide routes to genotypes that are either robust performers when challenged by environmental changes or high performers in more defined environments. In addition, G-by-G-by-E might provides multiple routes for populations and individuals to adapt to environmental changes. With the presence of environmentally independent additive effects, it is likely that alleles can be accumulated to intermediate frequencies in populations with small or no fitness costs in many environments. Upon rapid environmental change, the G-by-G-by-E interactions studied here can suppress or release large amounts of selectable genetic variation at a considerably higher rate^24–26^, facilitating more rapid and larger selection responses beyond predictions obtained based on the levels of additive variance, or heritabilities, in the populations.

### Connections to available knowledge

Our reanalyses of experimental data show that G-by-G-by-E interactions were generally important for the variation in growth across environments. This extends and connects earlier results from numerous studies in multiple species illustrating the importance of G-by-G and G-by-E for complex trait variation^1–7^. For example, many yeast genes are known to be nonessential in one genetic background, but essential in another, with the essential genes often being highly connected hubs in interaction networks^19^. Similar mechanisms have been studied also in bacteria where, for example, evaluations of the effects of 18 randomly selected mutations in *E.coli* in two environments and five genetic backgrounds illustrate that all of them have genetic background dependent effect on phenotypic plasticity^13^. Capacitation is also a well known mechanisms studied in detail in several species, including the heat shock protein *HSP90*^27,28^ in *Arabidopsis thaliana* and *EGFR* in Drosophila melanogaster^29^. Studying the connection between G-by-G and G-by-E requires large and powerful populations. Yeast is a model where such data can be generated and using large segregating experimental populations, single trait G-by-G-by-E interactions have been dissected in detail^19^. Our work connects and generalises these earlier findings from multiple species, populations and traits. We show how the network topology, and its underlying interactions, are likely central for the expression of complex phenotypes in individuals, and also how these effects are captured by quantitative genetics models^1^. Further we also illustrate how dynamic genetic effects are in response to environmental perturbations and the likely central role of genetic interactions in general and capacitating epistasis in particular for this. Despite the experimental and analytical challenges, these results highlight the importance of considering not only G-by-G and G-by-E, but also their mutual dependence on one another, when interpreting the results from complex trait genetic studies aiming to understand complex trait variation and evolution.

### Hypothetical molecular mechanisms underlying the observed effects

A number of the mapped hub-QTL harbour candidate genes with known biological functions, including *GPA1*^30^*, HAP1*^31–33^*, KRE33*^34^*, MKT1*^35^ and *IRA2*^36^ (Figure S3). Although they are obvious positional, further work is required to validate the functional candidate genes, in particular if and how they might contribute to the dynamic changes across environments. Not only because earlier work focussed on marginal genetic effects, rather than network effects, but also as the studies were performed in the environments where the effects were most prominent. Some earlier findings, however, allows us to present hypotheses about ways that they might contribute to the effects discovered here. For example, several hub-QTL were also epistatic hubs for expression QTLs^22^ where interactions between *HAP1* - *KRE33* and between *HAP1* - *MKT1* contributed to variations in the expression levels of many genes^22^. These studies were, however, performed in a single environment and expression QTL are known to often be environmentally dependent^37,38^. Further studies of transcriptomic and metabolic data across multiple environments would therefore be a possible route to explore whether changes in the topology of the genetic networks around these loci across environments results in associated changes also at transcriptomic and metabolic levels.

### Potential implications for modelling of quantitative traits

The studies of this yeast population here, and earlier^1^, illustrate that the most highly connected loci in the interaction networks (the hubs) often serve as modulators. They have little, or no, individual effects but rather influence the phenotype by releasing the effects of environmentally specific sets of interacting effector genes. This is an opposite scenario to that assumed in the recently proposed Omnigenetic model for quantitative traits^39^, where it is postulated that the highly connected loci in the networks are effectors that are modulated by many other genes. One consequence of this is that results from association studies where loci are detected based on their marginal additive effects need to be interpreted with caution. This is because it might be incorrect to assume that such effects suggest that the locus has a direct (effector) influence on the trait or disease, while it in fact could be entirely a composite effect of contributions by multiple other effector loci. Another potential modelling challenge highlighted here is the potentially large influence by epistasis by environment interactions. We find that they might not only influence the variation in quantitative traits by modulating the effects of individual genes, but also in defining which sets of interactors that are under genetic control by capacitor loci. Further theoretical, and empirical, work is needed to explore the potential implications of these findings for modelling of quantitative trait variation from molecular data in, for example, genome-wide association studies and studies on the basis for, maintenance of and utilization of genetic variation in short- and long-term adaptations to natural and artificial selection.

### Conclusions

In summary, we show that epistatic network rewiring is a common response to environmental perturbations. The dynamics in the network topology across environments were connected to changes in allelic effects of individual loci, epistatic effects of multi-locus interactions and the genetic variance contributed by these on the population level. These findings illustrated how G-by-G-by-E interactions influences both individual phenotypes and population level genetic variation. Our results provide novel insights on the fundamental mechanisms contributing to variation in complex traits with practical implications to, in particular, fields where the genetic mechanisms facilitating responses to variations in the environment are central, including evolutionary biology and breeding.

## Methods

### Downloaded data for the BY x RM haploid segregant yeast population

A detailed description of the generation of the BY × RM strains, as well as the genotyping, phenotyping, quality control of genotypes, filtering and normalization of growth measurements is available in Bloom *et al*^21^. All the data analysed here was downloaded from the supplements of that paper. The previously detected additive and epistatic QTLs, as well as their connectivity into within-environment interaction networks, are available in the supplementary information of Forsberg *et al*^1^.

### Estimating the contributions by genotype and genotype-by-environment interactions to the phenotypic variance

The phenotypic variance was partitioned into contributions from genotype (G), genotype-by-environment (G-by-E) and residual (environmental; E) effects by fitting model (1) to the data:

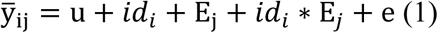

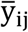 is the mean growth for the replicates of individual *i* in environment *j* (*j* = 1..*n*; *n* is the number of growth conditions); *id_i_* is the individual segregant (genotype) coded as a factor and *E_j_* is a dummy variable representing the growth condition (environment). *id_i_*∗*E_j_* is the interaction (G-by-E) between a particular segregant (genotype) and growth condition (environment). Since the available data was normalised against a control medium, there was (as expected) no significant contrition by *E*. The relative contributions to the total growth (phenotypic) variance from G and G-by-E were estimated by their respective sum of squares (Sum of Square for *id* is calculated as 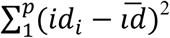 and Sum of Square for the interaction *id*∗*E* is calculated as 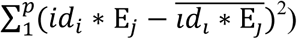

### Defining a set of independently associated additive growth loci

A set of across-environment growth loci was defined. First, QTLs detected in the earlier environment-separate analyses with peak associations within 20kb and in pairwise r^2^ > 0.9 were selected. Second, all the loci selected in step 1 were subjected to a multi-locus polygenic association analysis ^40,41^ to identify a final set of statistically independent loci (FDR < 0.05) with additive effects on growth in each tested environment^21^. Alternative definitions ranging from physical distance < 20 kb and r^2^ > 0.6 to physical distance < 10 kb and r^2^ > 0.9 were evaluated and found to result in very similar results in practice (result not shown).

### Across environment evaluation of the additive growth loci

Several growth loci in the final set defined above only had significant individual associations in one growth environment. To test if they, as a group, contributed to the polygenic inheritance of growth also in other environments we compared the fit of the following models to the data (models 2 and 3) using a likelihood ratio test.

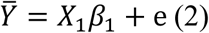

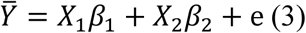

Here, 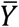 is a vector of the average growth of each segregant (genotype) in a particular environment and *e* is the norsmally distributed residual. The joint contributions by the individually significant/non-significant loci in a specific environment was modelled in *X*_1_*β*_1_/*X*_2_*β*_2_, respectively. *X_1_* includes a column vector of 1’s for the population mean and column vectors with the genotype of each significantly associated SNP in the environment with the two homozygous genotypes coded as 0/2, respectively. *X*_2_ includes column vectors with genotypes of all loci in the set defined above that was not individually significant in the tested environment. *β*_1_/*β*_2_ are vectors including the estimated additive effects for the two sets of loci. A likelihood ratio test was used to compare the fit of the two models using the *lrtest* function in R package *lmtest*^42^.

### Detect individual loci involved in genotype by environment interactions

All growth loci defined in the polygenic analysis were evaluated for genotype by environment interactions. This by fitting the following two models to the data:

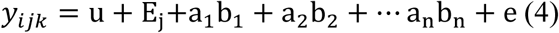

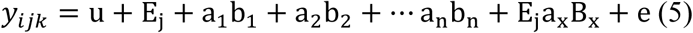

In both models, *y_ijk_* is the growth of replicate *k* for segregant *i* in environment *j* (*j* = 1..20 environments; *k* = 1..*n_ij_*; *n_ij_* is the number of replicates for individual *i* in environment *j*); *a_x_* is the indicator regression variable for the genotype of QTL *x* coded as 0 and 2 for the homozygous minor and major alleles; *b_x_* are the corresponding estimated additive effects; *u* is the population mean and *E_j_* is the effect of environment *j* (j = 1…20) on growth. Model 5 also includes an interaction term E_j_a_x_B_x_ between one of the QTL and the environment. Model 5 was fitted for each QTL one at a time to test for its interaction with the 20 growth environments. The significance of each QTL by environment interaction was evaluated using a likelihood ratio test between models 4 and 5. Polygenicity was accounted for by the simultaneous fitting of all mapped loci in the two models. The analyses were performed using custom R scripts^43^.

### Estimating the additive effects of QTL in different growth environments

A linear model (model 6) was used to estimate the additive effects of all the additive loci selected in the polygenic analysis in each tested environment.

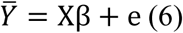

Here, 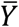 is the average growth of the replicates for each segregant (genotype) in each tested condition. *e* is the normally distributed residual. *X* includes a column vector of 1’s for the population mean and additional column vectors with the indicator regression variables for all the SNPs included in the model (coded as 0/2 for the two homozygous genotypes, respectively). β is a column vector with the corresponding additive effects. This model was fitted for each environment independently to obtain estimates of the additive effects for each locus in each environment.

### Construction of an across-environment epistatic interaction network

A complete across-environment epistatic growth interaction network was constructed from the environment specific networks reported in Forsberg *et al*^1^. First, the environment specific networks inferred for each growth environment were extracted from the results of *Forsberg et al*^1^. Second, we evaluated whether any of the pairwise interactions detected for a specific environment made significant contribution to growth at the remaining environments under a more lenient significant threshold only correcting for the total number of pairwise interactions detected across all environments. This was performed using a likelihood ratio test between models with and without the pairwise interaction for a particular pair of epistatic loci as described in detail by *Bloom et al*^21^. Then, the across-environment network was constructed by connecting the loci display pairwise interaction in any of the 20 environments using *igraph*^44^ as descried in *Forsberg et al*^1^. These analyses were performed using custom scripts implemented in R^43^ [will be made available upon publication or by request during review]. The raw data from which these were constructed are avaiable as supplement 5 in Bloom et al ^21^; The analysis scripts/results from these earlier analyses are available at https://github.com/simfor/yeast-epistasis-paper and are described in detail by *Forsberg et al*^1^).

### Evaluation of the G-by-G-by-E interaction for a six-locus interaction network

The effects of a six locus interaction network, originally detected and explored for growth in IAA containing growth medium in Forsberg *et al*^1^, were here evaluated across multiple environments. This analysis was performed to explore the association between the rewiring of the epistatic genetic network and its contribution to growth in the different environments. To quantify the effects of G-by-G-by-E interactions, models 4 and 5 were fitted to the data with a_x_ (*x* = *n* =1) used as the indicator variable for each of the 64-genotype classes defined by the genotypes at the 6 loci. All other parameters, and the likelihood ratio test used to obtain the P-values for comparing the models, were the same as described above.

## List of abbreviation

G-by-G interaction: Genotype by Genotype interaction

G-by-E interaction: Genotype by Environment interaction

G-by-G-by-E interaction: Genotype by Genotype by Environment interaction

## Acknowledgements

We thank Simon Forsberg, Tilman Rönneburg and Thibaut Payen for discussions and comments on the manuscripts and figures.

## Additional files

All supplementary figures and tables are available in Additional file1

## Authors’ contributions

ÖC and YZ initiated the study and designed the project; YZ performed the data analyses; ÖC and YZ summarised the results and wrote the manuscript. Both authors read and approved the final manuscript.

---

**Figure S1.**
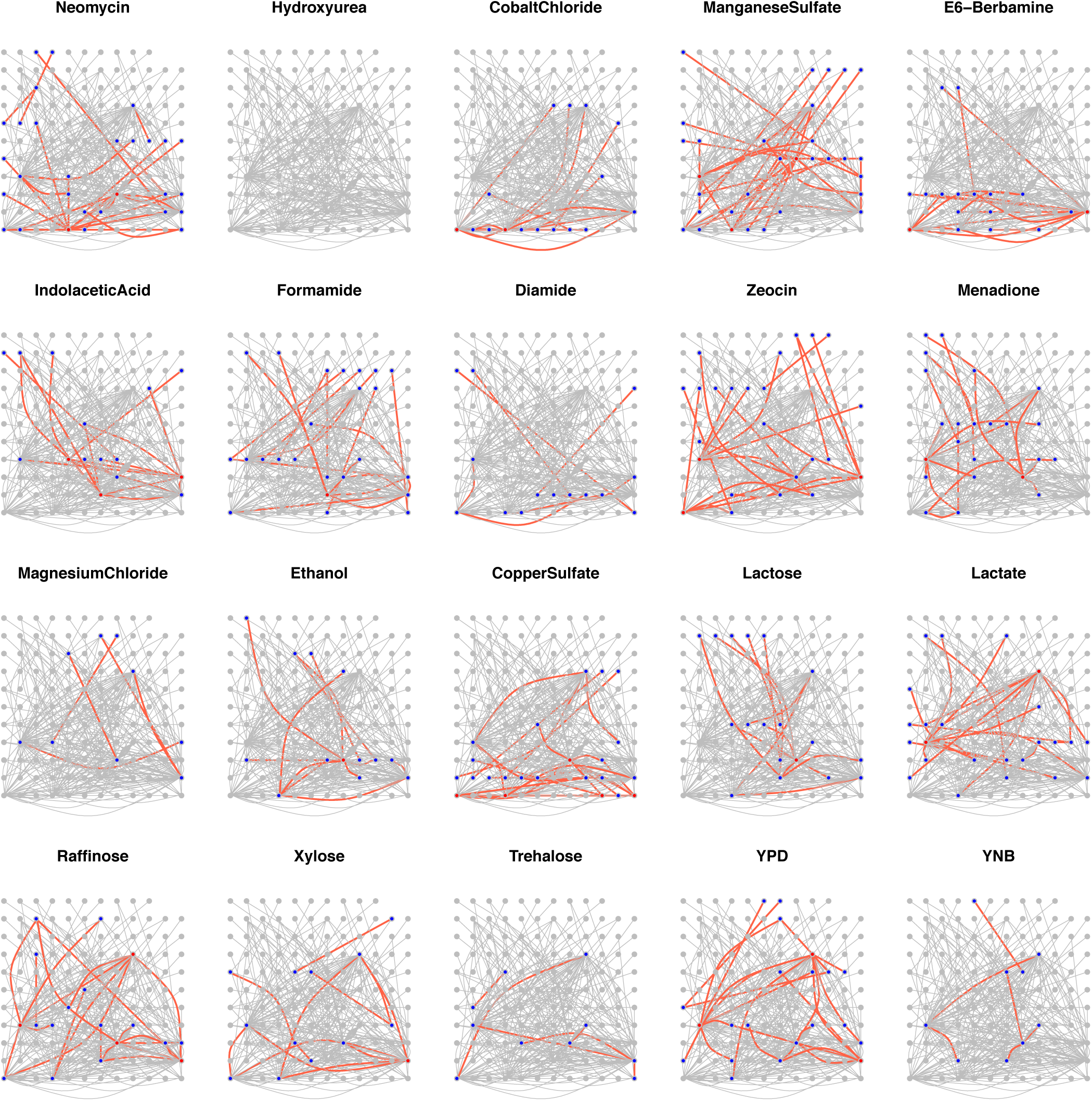
Joint epistatic network constructed by connecting shared loci from 20 epistatic networks detected for each media/environment. Each dot in this plot is a loci and the colour of the dot describes if the loci is detected with epistatic interaction for the current medium with yes being blue or red (connected with more than 4 other loci) and no being grey. The pairwise interactions between loci are indicted by connected edges. The number of edges connecting two loci describes the number of times it is detected across 20 mediums, and the detected connection for current medium is highlighted with red (detected in Bloom et al,), blue (our study) and grey (other medias).

**Figure S2.**
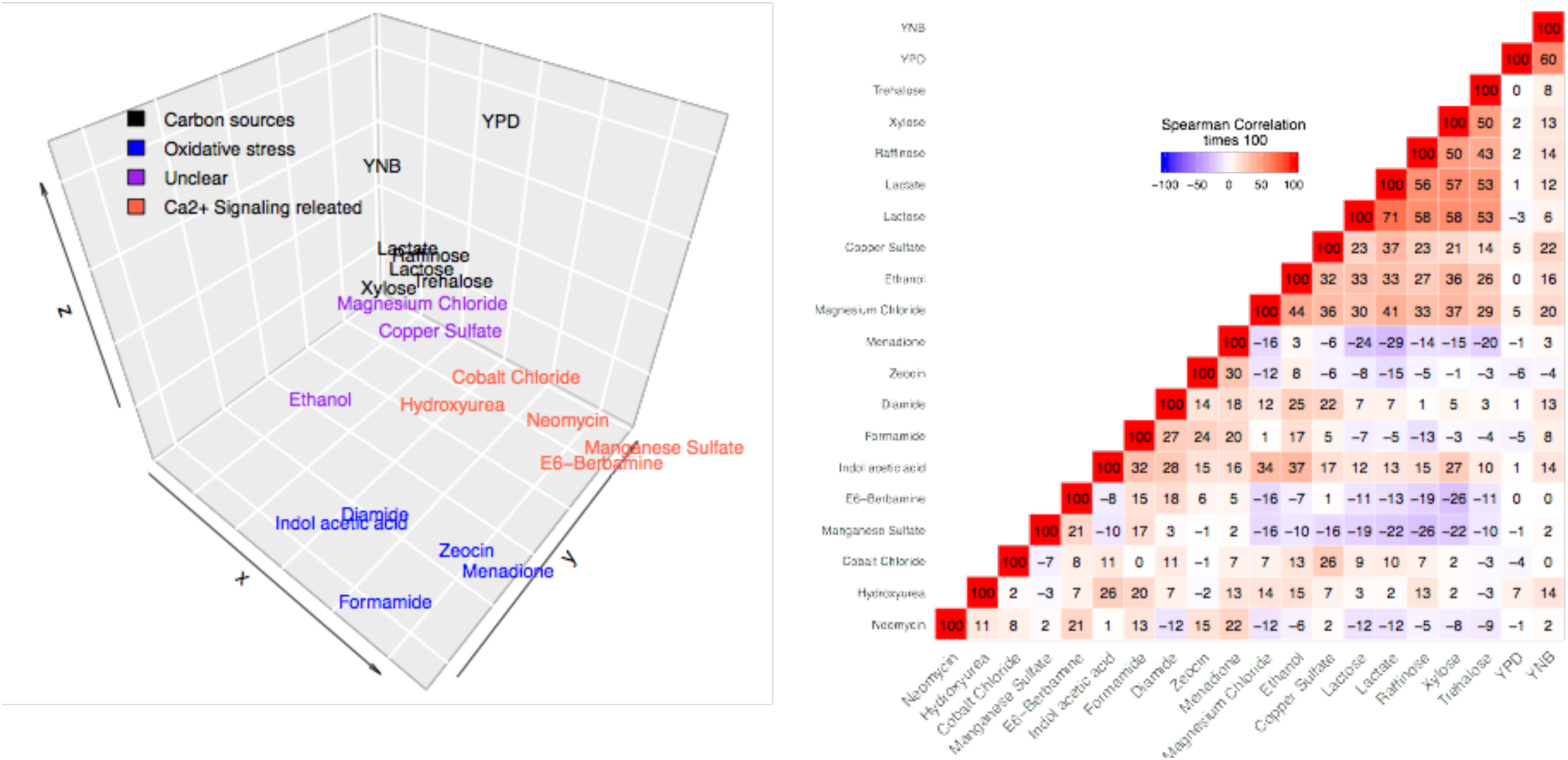
Illustration of the phenotype resemblance and change of phenotypic/genetic variance under different growth condition. **A**). 3-dimentainal PCA plot of the yeast growth measured as the radial of colony on 20 different mediums. These mediums were made by adding small-chemical molelues to mimic different enviroments ^11^. **B**). Pairwise Spearman rank correlation among growth measured on these 20 mediums. Numbers in the cell are 100 times the Spearman correation, and environments were sorted based on their order after hierarchical cluster.

**Figure S3.**
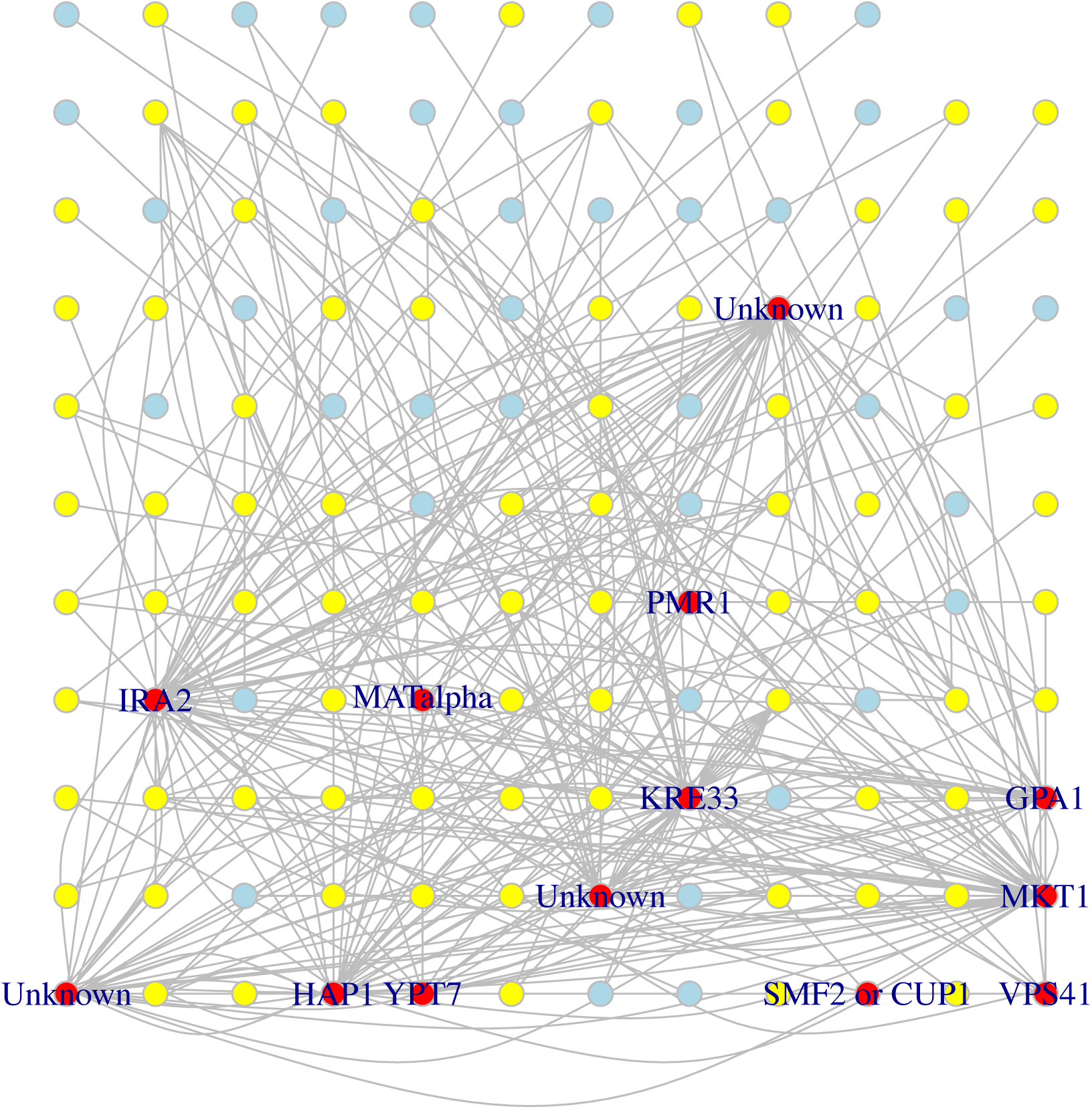
Illustration of the connectivity of the 13 hubs. 13 hubs connected with more than 4 loci in at least 1 environment is highlighted in red, loci epistatically interact with these hubs in at least one environment are labeled in yellow.

**Figure S4.**
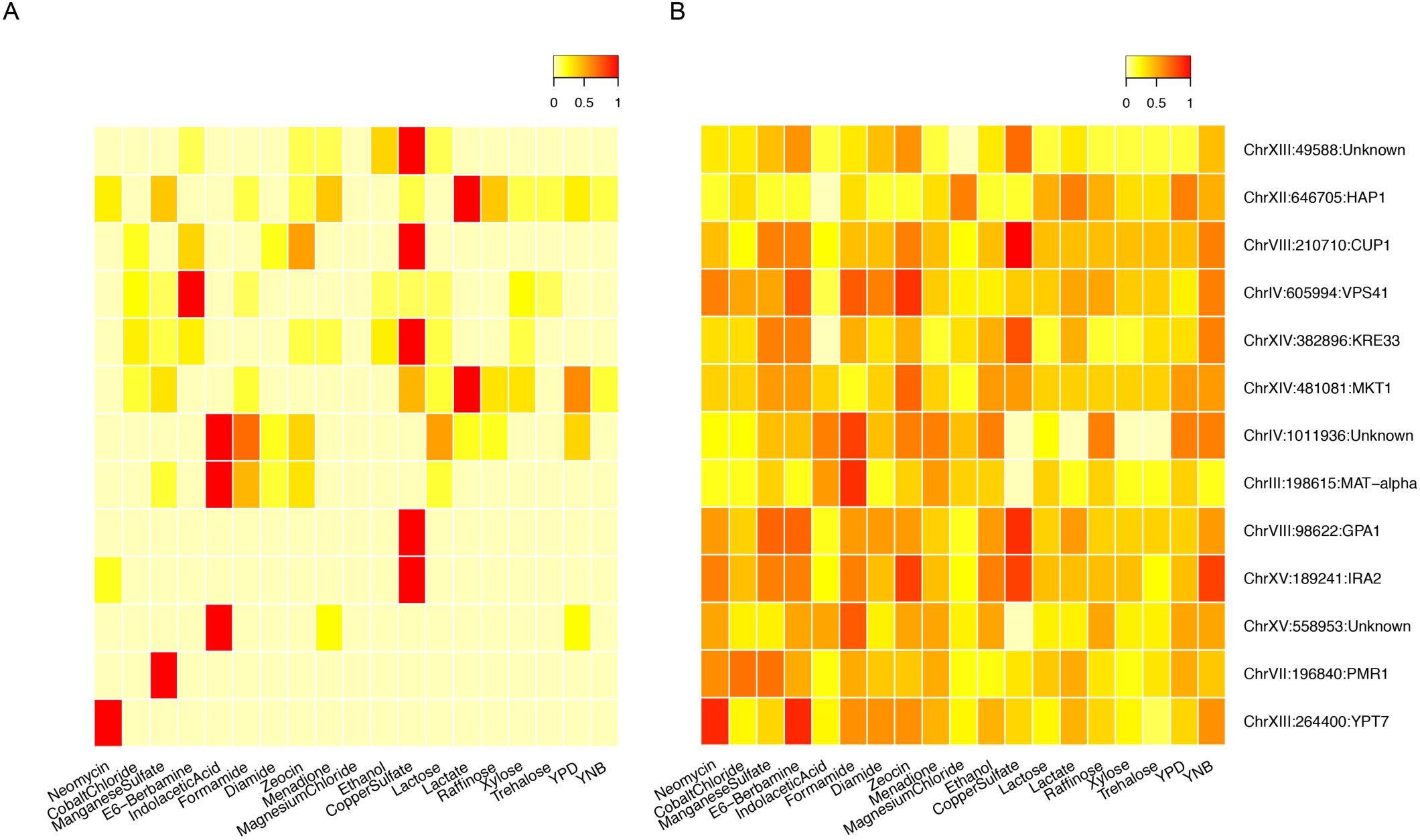
Rewiring of the 13 epistatic networks across 20 environments. **A)** Proportion of epistatically connected loci for a specific environment. Each row of the heatmap represents a network with the hub alleles and candidate genes marked to the left and each column represents an environment. The environments are sorted based on their order after hierarchical cluster. **B)** Histogram of the proportion of deactivated loci, including these either remained either as additive loci or rewired to other loci.

**Figure S5.**
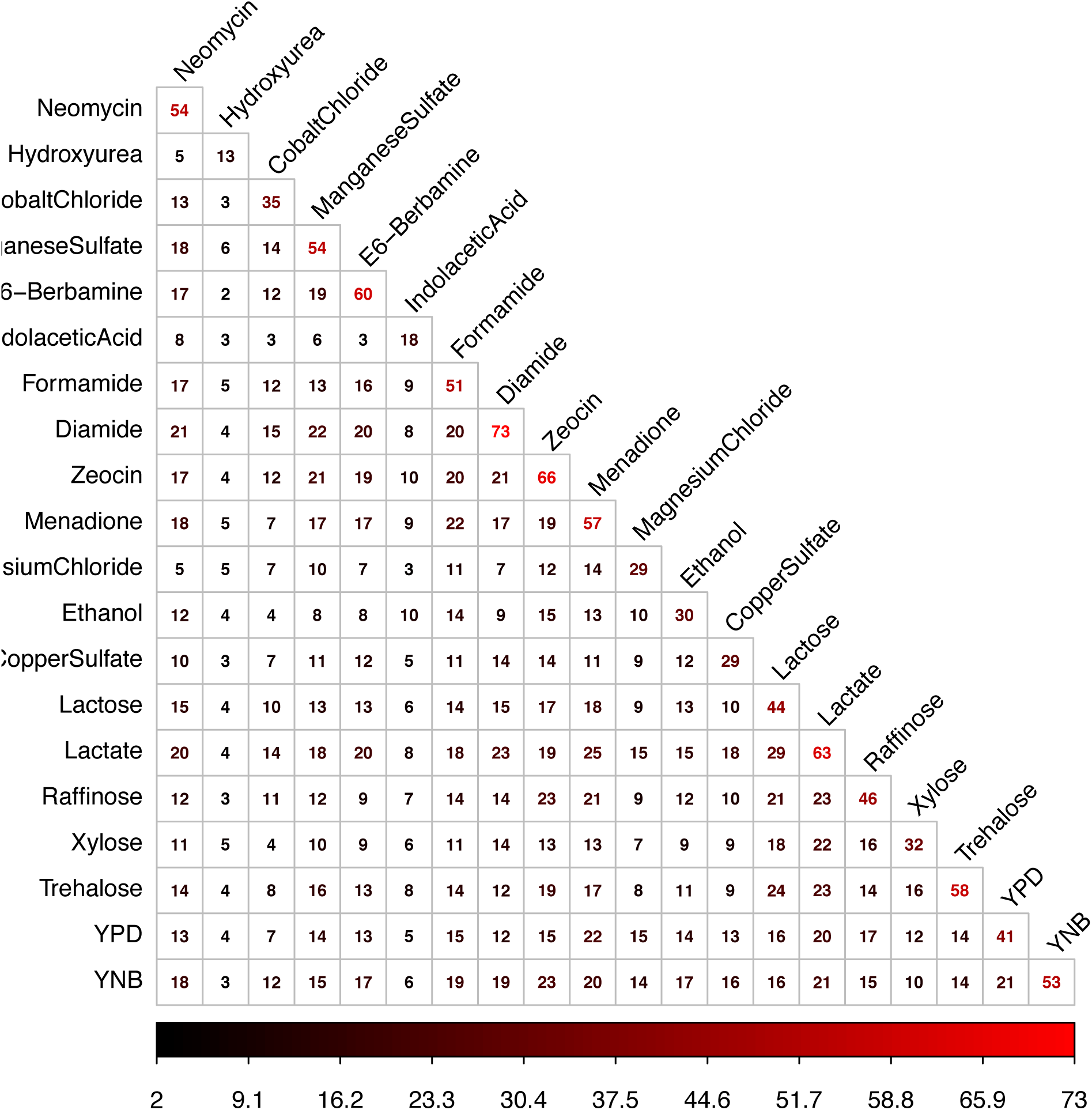
Pairwise overlap of loci detected with additive effects. Numbers on the diagnal is the number of addtive loci detected for a particular enviroment, and numbers in the cell are the number of overlap addtive loci. Phenotype were sorted based on their order after hierarchical cluster.

**Figure S6.**
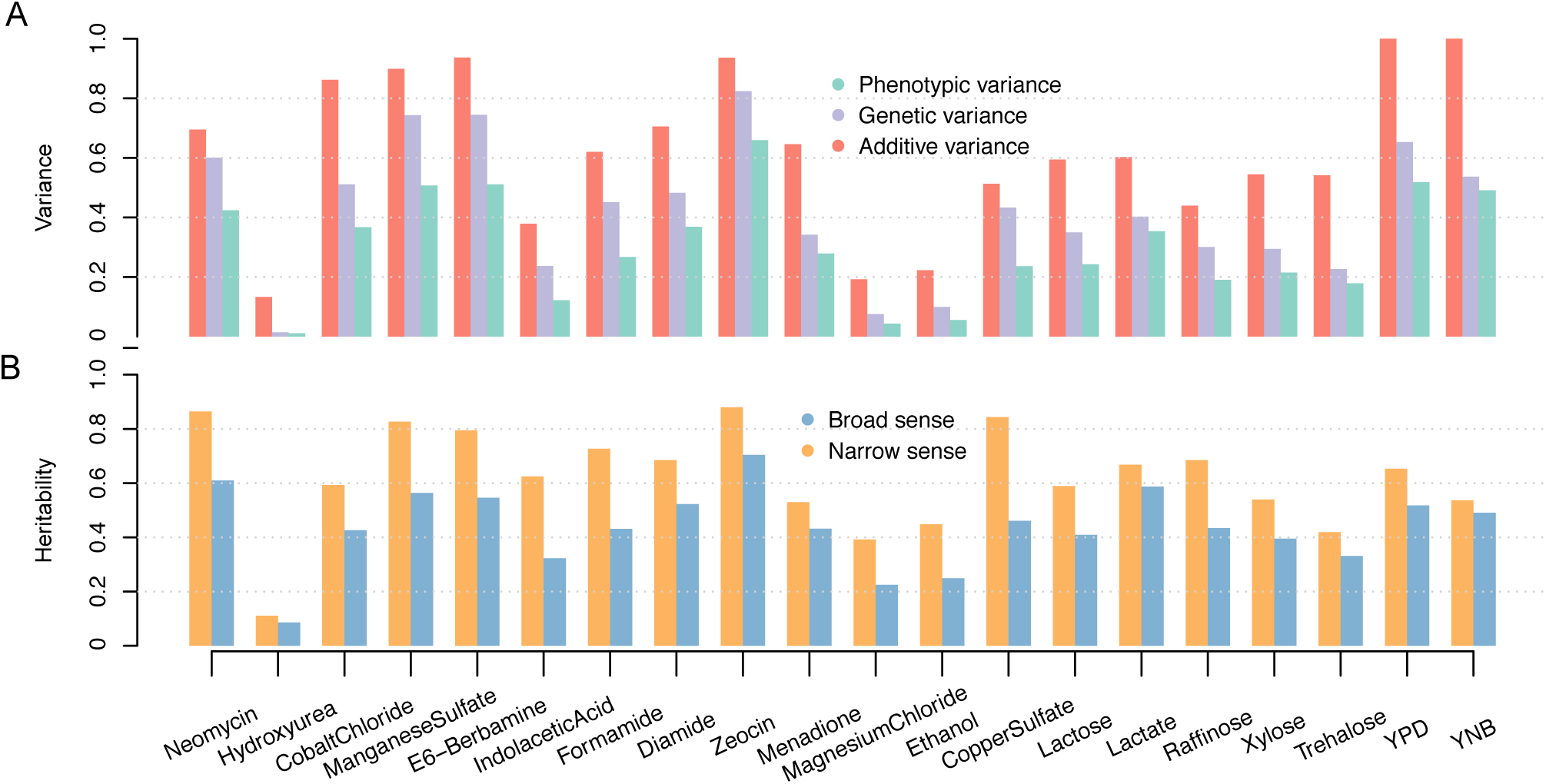
Change of genetic variance and broad/narrow sense heritability. **A)**. The phenotypic variance, genetic variance and additive variance for growth under 20 midums. Genetic variance and additive variance was estimated as the product of phentypic variance and broad- and narrow-sense heritability (panel **B**) reported in Bloom et al ^1^.

**Figure S7.**
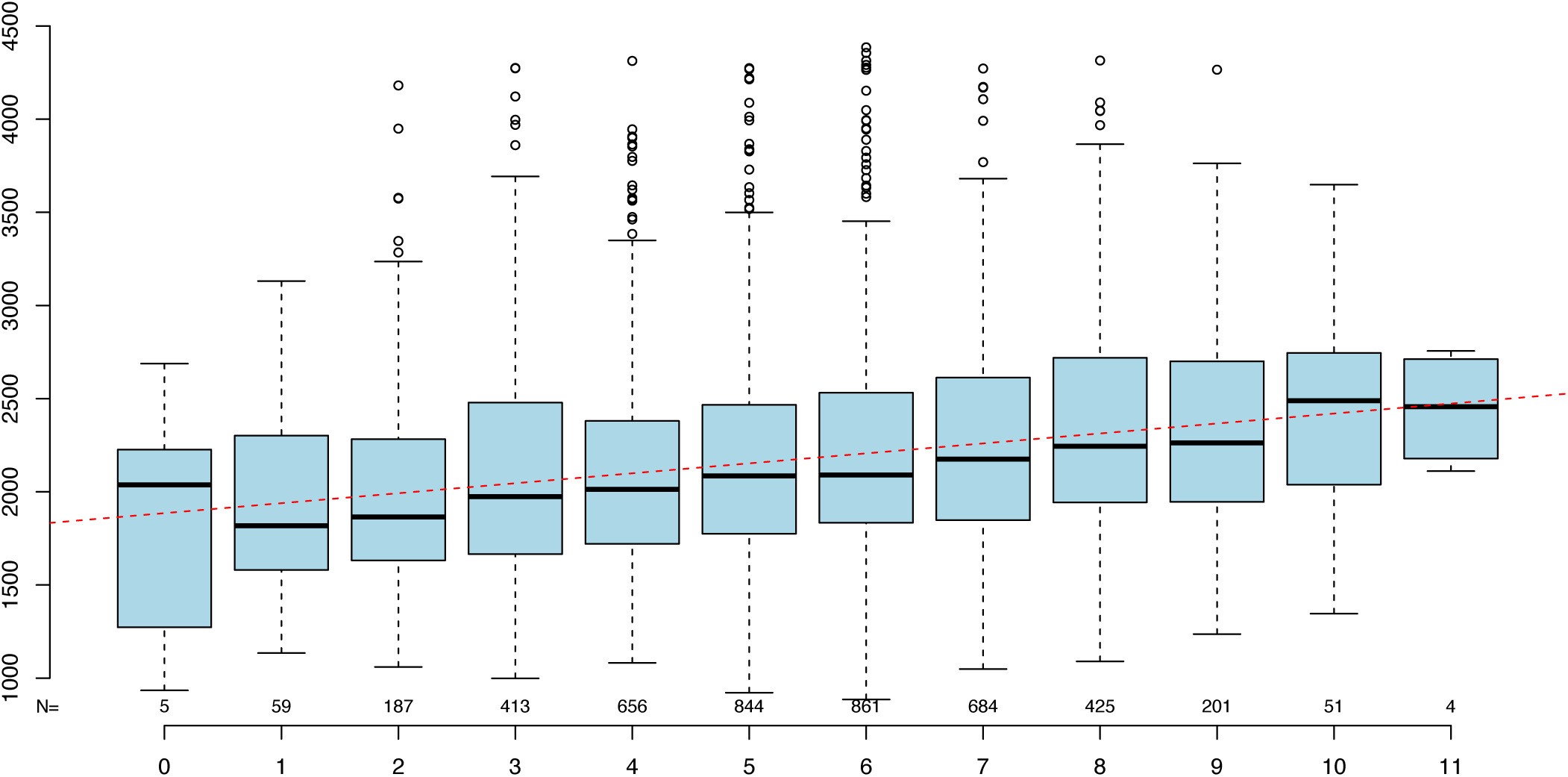
Releationship between the mean growth rank across 20 enviroments and the number of non-conpacitated alleles. X-axis is the number of non-compacitated alleles across 13 hubs detected in our study, and y-axis is the mean growth rank obtained by first rank the growth measurements across 20 enviroemtns and then taking the artihma tical mean.

**Table S1.** Summary of the P values fron likelyhood ratio test comparing a full model with all 311 growth QTL detected for all enviroments and a reduced model with QTL only detected for focal enviroments

| Environment | Number of QTL<sup>1</sup> | P values<sup>2</sup> | Significance<sup>3</sup> |
| --- | --- | --- | --- |
| CobaltChloride | 35 | 3.44e-04 | Yes |
| CopperSulfate | 29 | 8.44e-04 | Yes |
| Diamide | 73 | 2.27e-04 | Yes |
| E6-Berbamine | 60 | 1.30e-03 | Yes |
| Ethanol | 30 | 4.35e-05 | Yes |
| Formamide | 51 | 4.64e-03 | No |
| Hydroxyurea | 13 | 4.03e-04 | Yes |
| IndolaceticAcid | 18 | 2.56e-04 | Yes |
| Lactate | 63 | 2.89e-03 | No |
| Lactose | 44 | 1.06e-03 | Yes |
| MagnesiumChloride | 29 | 4.80e-04 | Yes |
| ManganeseSulfate | 54 | 8.38e-05 | Yes |
| Menadione | 57 | 1.09e-02 | No |
| Neomycin | 54 | 1.96e-11 | Yes |
| Raffinose | 46 | 1.06e-05 | Yes |
| Trehalose | 58 | 5.71e-03 | No |
| Xylose | 32 | 1.58e-09 | Yes |
| YNB | 53 | 1.06e-04 | Yes |
| YPD | 41 | 1.04e-04 | Yes |
| Zeocin | 66 | 3.43e-06 | Yes |

<sup>1</sup>Number of QTL: The number of additive QTL detected at focal environment
<sup>2</sup>P values from likelihood ratio test testing whether these loci, not individually associated with focal environments made significant contribution to growth as a group
<sup>3</sup>Significance: whether the p value reach multiple testing

**Table S2.** Summary of the QTL by E analysis for 311 growth QTL

| SNP Name | Chromosome | Position | P_vlaue | GxE<sup>1</sup> |
| --- | --- | --- | --- | --- |
| 193500_chrI_193500_C_T | 1 | 193500 | 3.88e-242 | Yes |
| 4348634_chrVII_609644_T_C | 7 | 609644 | 3.62e-84 | Yes |
| 6164100_chrX_331639_C_T | 10 | 331639 | 9.88e-12 | Yes |
| 3089307_chrV_197352_T_C | 5 | 197352 | 5.32e-58 | Yes |
| 8347770_chrXIII_24565_A_G | 13 | 24565 | 0.00e+00 | Yes |
| 6493214_chrX_660753_C_T | 10 | 660753 | 6.16e-127 | Yes |
| 7894865_chrXII_649837_A_G | 12 | 649837 | 0.00e+00 | Yes |
| 10107829_chrXV_75860_A_G | 15 | 75860 | 5.95e-300 | Yes |
| 4111213_chrVII_372223_T_G | 7 | 372223 | 1.28e-73 | Yes |
| 9734886_chrXIV_487250_G_C | 14 | 487250 | 0.00e+00 | Yes |
| 10590922_chrXV_558953_G_A | 15 | 558953 | 8.92e-140 | Yes |
| 3175379_chrV_283424_T_C | 5 | 283424 | 8.24e-68 | Yes |
| 1139178_chrIII_95776_T_C | 3 | 95776 | 2.42e-49 | Yes |
| 9238100_chrXIII_914895_A_T | 13 | 914895 | 3.13e-54 | Yes |
| 10973374_chrXV_941405_C_T | 15 | 941405 | 5.77e-19 | Yes |
| 7815060_chrXII_570032_G_A | 12 | 570032 | 1.48e-232 | Yes |
| 4951944_chrVIII_122014_T_C | 8 | 122014 | 0.00e+00 | Yes |
| 10233761_chrXV_201792_T_C | 15 | 201792 | 0.00e+00 | Yes |
| 40866_chrI_40866_T_C | 1 | 40866 | 5.18e-154 | Yes |
| 3661374_chrVI_192545_C_T | 6 | 192545 | 2.64e-08 | Yes |
| 7614510_chrXII_369482_A_G | 12 | 369482 | 1.74e-59 | Yes |
| 10875861_chrXV_843892_A_G | 15 | 843892 | 2.28e-30 | Yes |
| 7379379_chrXII_134351_T_C | 12 | 134351 | 2.76e-43 | Yes |
| 1255174_chrIII_211772_G_A | 3 | 211772 | 6.91e-31 | Yes |
| 796888_chrII_566670_T_A | 2 | 566670 | 7.23e-88 | Yes |
| 9790206_chrXIV_542570_G_A | 14 | 542570 | 3.72e-235 | Yes |
| 8372793_chrXIII_49588_C_T | 13 | 49588 | 0.00e+00 | Yes |
| 10465962_chrXV_433993_C_T | 15 | 433993 | 1.90e-127 | Yes |
| 8434826_chrXIII_111621_T_C | 13 | 111621 | 0.00e+00 | Yes |
| 2308435_chrIV_948413_A_G | 4 | 948413 | 6.39e-101 | Yes |
| 421432_chrII_191214_G_A | 2 | 191214 | 3.44e-23 | Yes |
| 5892760_chrX_60299_C_T | 10 | 60299 | 6.23e-89 | Yes |
| 10170631_chrXV_138662_A_G | 15 | 138662 | 0.00e+00 | Yes |
| 11644024_chrXVI_520764_G_A | 16 | 520764 | 7.39e-102 | Yes |
| 5572582_chrIX_180009_T_A | 9 | 180009 | 6.19e-38 | Yes |
| 10196253_chrXV_164284_T_A | 15 | 164284 | 0.00e+00 | Yes |
| 5441452_chrIX_48879_C_T | 9 | 48879 | 3.59e-25 | Yes |
| 7193645_chrXI_615433_T_C | 11 | 615433 | 2.10e-62 | Yes |
| 1974760_chrIV_614738_A_C | 4 | 614738 | 4.04e-36 | Yes |
| 5040640_chrVIII_210710_A_T | 8 | 210710 | 4.06e-122 | Yes |
| 3918900_chrVII_179910_G_A | 7 | 179910 | 0.00e+00 | Yes |
| 4590187_chrVII_851197_T_G | 7 | 851197 | 2.32e-70 | Yes |
| 9624126_chrXIV_376490_A_G | 14 | 376490 | 0.00e+00 | Yes |
| 11615614_chrXVI_492354_C_T | 16 | 492354 | 2.61e-118 | Yes |
| 9142223_chrXIII_819018_C_T | 13 | 819018 | 1.20e-40 | Yes |
| 774352_chrII_544134_G_A | 2 | 544134 | 7.47e-111 | Yes |
| 623487_chrII_393269_T_C | 2 | 393269 | 3.42e-96 | Yes |
| 3866036_chrVII_127046_A_G | 7 | 127046 | 0.00e+00 | Yes |
| 3153148_chrV_261193_A_T | 5 | 261193 | 1.46e-52 | Yes |
| 2834907_chrIV_1474885_A_G | 4 | 1474885 | 2.89e-07 | Yes |
| 8560038_chrXIII_236833_C_T | 13 | 236833 | 0.00e+00 | Yes |
| 10067884_chrXV_35915_T_C | 15 | 35915 | 2.00e-93 | Yes |
| 1295281_chrIII_251879_C_T | 3 | 251879 | 1.58e-16 | Yes |
| 9711190_chrXIV_463554_A_T | 14 | 463554 | 0.00e+00 | Yes |
| 5011062_chrVIII_181132_A_G | 8 | 181132 | 2.17e-105 | Yes |
| 5531209_chrIX_138636_T_A | 9 | 138636 | 3.78e-57 | Yes |
| 8231374_chrXII_986346_G_A | 12 | 986346 | 1.01e-08 | Yes |
| 5581492_chrIX_188919_A_T | 9 | 188919 | 6.04e-51 | Yes |
| 5666111_chrIX_273538_G_A | 9 | 273538 | 1.22e-78 | Yes |
| 11498533_chrXVI_375273_C_T | 16 | 375273 | 2.23e-146 | Yes |
| 3701448_chrVI_232619_G_A | 6 | 232619 | 2.39e-07 | Yes |
| 487143_chrII_256925_T_C | 2 | 256925 | 5.88e-45 | Yes |
| 4945907_chrVIII_115977_A_T | 8 | 115977 | 0.00e+00 | Yes |
| 684479_chrII_454261_G_A | 2 | 454261 | 1.52e-133 | Yes |
| 1118075_chrIII_74673_A_G | 3 | 74673 | 3.48e-33 | Yes |
| 4537389_chrVII_798399_T_C | 7 | 798399 | 1.36e-56 | Yes |
| 8587605_chrXIII_264400_A_G | 13 | 264400 | 0.00e+00 | Yes |
| 10280547_chrXV_248578_G_A | 15 | 248578 | 9.41e-211 | Yes |
| 3936455_chrVII_197465_A_G | 7 | 197465 | 0.00e+00 | Yes |
| 10653941_chrXV_621972_A_G | 15 | 621972 | 8.37e-84 | Yes |
| 4216167_chrVII_477177_C_G | 7 | 477177 | 1.23e-119 | Yes |
| 9750134_chrXIV_502498_C_T | 14 | 502498 | 0.00e+00 | Yes |
| 10688536_chrXV_656567_G_A | 15 | 656567 | 2.85e-58 | Yes |
| 884725_chrII_654507_T_C | 2 | 654507 | 1.19e-47 | Yes |
| 4139152_chrVII_400162_T_C | 7 | 400162 | 4.15e-76 | Yes |
| 5290246_chrVIII_460316_T_C | 8 | 460316 | 4.70e-30 | Yes |
| 6787397_chrXI_209185_T_A | 11 | 209185 | 2.78e-44 | Yes |
| 5653342_chrIX_260769_T_C | 9 | 260769 | 3.51e-125 | Yes |
| 5430557_chrIX_37984_T_G | 9 | 37984 | 8.27e-23 | Yes |
| 9385314_chrXIV_137678_C_T | 14 | 137678 | 5.60e-33 | Yes |
| 7534566_chrXII_289538_G_A | 12 | 289538 | 8.28e-32 | Yes |
| 3823704_chrVII_84714_T_A | 7 | 84714 | 2.53e-164 | Yes |
| 8073782_chrXII_828754_T_C | 12 | 828754 | 6.92e-76 | Yes |
| 4994073_chrVIII_164143_G_A | 8 | 164143 | 1.73e-109 | Yes |
| 10442033_chrXV_410064_A_G | 15 | 410064 | 5.13e-175 | Yes |
| 7298297_chrXII_53269_C_T | 12 | 53269 | 3.91e-09 | Yes |
| 7693396_chrXII_448368_A_G | 12 | 448368 | 9.07e-33 | Yes |
| 10134640_chrXV_102671_A_T | 15 | 102671 | 0.00e+00 | Yes |
| 8736983_chrXIII_413778_C_T | 13 | 413778 | 2.67e-76 | Yes |
| 6354106_chrX_521645_T_G | 10 | 521645 | 1.04e-101 | Yes |
| 11541717_chrXVI_418457_T_C | 16 | 418457 | 1.49e-100 | Yes |
| 1641349_chrIV_281327_T_C | 4 | 281327 | 1.82e-38 | Yes |
| 2717811_chrIV_1357789_T_G | 4 | 1357789 | 8.57e-27 | Yes |
| 8654539_chrXIII_331334_A_G | 13 | 331334 | 1.25e-276 | Yes |
| 7110331_chrXI_532119_G_A | 11 | 532119 | 1.94e-37 | Yes |
| 1902084_chrIV_542062_T_C | 4 | 542062 | 3.40e-35 | Yes |
| 2630055_chrIV_1270033_C_T | 4 | 1270033 | 2.43e-12 | Yes |
| 9512173_chrXIV_264537_A_G | 14 | 264537 | 9.43e-173 | Yes |
| 4299073_chrVII_560083_G_A | 7 | 560083 | 2.07e-109 | Yes |
| 10615684_chrXV_583715_A_G | 15 | 583715 | 8.44e-112 | Yes |
| 2509052_chrIV_1149030_T_C | 4 | 1149030 | 9.65e-56 | Yes |
| 33147_chrl_33147_G_T | 1 | 33147 | 7.86e-130 | Yes |
| 10858985_chrXV_827016_A_T | 15 | 827016 | 5.50e-23 | Yes |
| 9184667_chrXIII_861462_G_A | 13 | 861462 | 4.51e-55 | Yes |
| 8664927_chrXIII_341722_T_C | 13 | 341722 | 1.87e-227 | Yes |
| 3599017_chrVI_130188_C_T | 6 | 130188 | 4.33e-27 | Yes |
| 4397128_chrVII_658138_T_C | 7 | 658138 | 7.50e-22 | Yes |
| 7767974_chrXII_522946_A_G | 12 | 522946 | 1.75e-58 | Yes |
| 6887963_chrXI_309751_T_G | 11 | 309751 | 8.70e-34 | Yes |
| 6970318_chrXI_392106_C_T | 11 | 392106 | 9.65e-51 | Yes |
| 10798650_chrXV_766681_G_A | 15 | 766681 | 6.53e-07 | Yes |
| 1291951_chrIII_248549_C_T | 3 | 248549 | 5.10e-14 | Yes |
| 5492996_chrIX_100423_T_C | 9 | 100423 | 3.32e-59 | Yes |
| 4161376_chrVII_422386_C_T | 7 | 422386 | 4.99e-78 | Yes |
| 6340986_chrX_508525_G_A | 10 | 508525 | 5.98e-95 | Yes |
| 3560948_chrVI_92119_A_G | 6 | 92119 | 4.45e-18 | Yes |
| 9084976_chrXIII_761771_C_T | 13 | 761771 | 8.68e-42 | Yes |
| 8260775_chrXII_1015747_G_A | 12 | 1015747 | 3.57e-03 | No |
| 114652_chrl_114652_C_T | 1 | 114652 | 4.26e-28 | Yes |
| 9897446_chrXIV_649810_A_G | 14 | 649810 | 2.91e-117 | Yes |
| 11330697_chrXVI_207437_T_A | 16 | 207437 | 6.98e-89 | Yes |
| 1782092_chrIV_422070_C_T | 4 | 422070 | 4.34e-43 | Yes |
| 5881260_chrX_48799_G_A | 10 | 48799 | 1.34e-63 | Yes |
| 665194_chrII_434976_G_T | 2 | 434976 | 2.63e-121 | Yes |
| 1089964_chrIII_46562_T_A | 3 | 46562 | 1.28e-09 | Yes |
| 11936407_chrXVI_813147_T_C | 16 | 813147 | 2.05e-06 | Yes |
| 1422346_chrIV_62324_G_C | 4 | 62324 | 8.79e-21 | Yes |
| 5202635_chrVIII_372705_C_A | 8 | 372705 | 2.88e-21 | Yes |
| 4251340_chrVII_512350_A_T | 7 | 512350 | 3.59e-119 | Yes |
| 6225749_chrX_393288_T_C | 10 | 393288 | 2.10e-33 | Yes |
| 3676508_chrVI_207679_A_C | 6 | 207679 | 1.17e-08 | Yes |
| 964804_chrII_734586_C_T | 2 | 734586 | 2.19e-16 | Yes |
| 8178548_chrXII_933520_G_T | 12 | 933520 | 8.70e-27 | Yes |
| 7356451_chrXII_111423_A_G | 12 | 111423 | 4.21e-37 | Yes |
| 5634465_chrIX_241892_A_G | 9 | 241892 | 5.25e-71 | Yes |
| 3418667_chrV_526712_A_T | 5 | 526712 | 5.71e-11 | Yes |
| 9024358_chrXIII_701153_G_C | 13 | 701153 | 2.16e-136 | Yes |
| 3301125_chrV_409170_T_C | 5 | 409170 | 3.23e-39 | Yes |
| 7349496_chrXII_104468_A_T | 12 | 104468 | 6.43e-31 | Yes |
| 9961304_chrXIV_713668_T_C | 14 | 713668 | 2.39e-18 | Yes |
| 7504079_chrXII_259051_C_T | 12 | 259051 | 8.74e-20 | Yes |
| 2424307_chrIV_1064285_G_A | 4 | 1064285 | 1.02e-35 | Yes |
| 1521242_chrIV_161220_A_G | 4 | 161220 | 1.47e-15 | Yes |
| 8060554_chrXII_815526_A_G | 12 | 815526 | 2.58e-87 | Yes |
| 10341838_chrXV_309869_T_C | 15 | 309869 | 4.74e-183 | Yes |
| 3266770_chrV_374815_G_A | 5 | 374815 | 7.68e-46 | Yes |
| 3534631_chrVI_65802_T_G | 6 | 65802 | 2.53e-13 | Yes |
| 5348875_chrVIII_518945_A_G | 8 | 518945 | 7.82e-35 | Yes |
| 1220483_chrIII_177081_G_C | 3 | 177081 | 3.16e-50 | Yes |
| 9195907_chrXIII_872702_T_C | 13 | 872702 | 1.49e-58 | Yes |
| 10493762_chrXV_461793_G_A | 15 | 461793 | 1.01e-54 | Yes |
| 6806359_chrXI_228147_C_A | 11 | 228147 | 4.47e-38 | Yes |
| 1309673_chrIII_266271_G_A | 3 | 266271 | 1.96e-12 | Yes |
| 1999059_chrIV_639037_G_T | 4 | 639037 | 3.86e-22 | Yes |
| 6937354_chrXI_359142_T_C | 11 | 359142 | 8.99e-46 | Yes |
| 8733525_chrXIII_410320_T_C | 13 | 410320 | 5.42e-97 | Yes |
| 4684968_chrVII_945978_C_T | 7 | 945978 | 9.19e-32 | Yes |
| 8474614_chrXIII_151409_T_C | 13 | 151409 | 4.72e-292 | Yes |
| 5945399_chrX_112938_T_A | 10 | 112938 | 8.11e-240 | Yes |
| 1240708_chrIII_197306_T_C | 3 | 197306 | 1.24e-51 | Yes |
| 2561643_chrIV_1201621_C_T | 4 | 1201621 | 1.15e-37 | Yes |
| 5271763_chrVIII_441833_A_G | 8 | 441833 | 2.29e-35 | Yes |
| 2358650_chrIV_998628_A_T | 4 | 998628 | 2.95e-126 | Yes |
| 3396166_chrV_504211_A_C | 5 | 504211 | 1.04e-13 | Yes |
| 3342341_chrV_450386_C_T | 5 | 450386 | 3.57e-31 | Yes |
| 6713295_chrXI_135083_A_G | 11 | 135083 | 1.98e-52 | Yes |
| 10976399_chrXV_944430_T_C | 15 | 944430 | 2.90e-16 | Yes |
| 6625055_chrXI_46843_C_T | 11 | 46843 | 2.90e-43 | Yes |
| 8966073_chrXIII_642868_G_A | 13 | 642868 | 1.94e-86 | Yes |
| 3103942_chrV_211987_C_G | 5 | 211987 | 5.70e-50 | Yes |
| 4039180_chrVII_300190_G_A | 7 | 300190 | 4.64e-87 | Yes |
| 7473402_chrXII_228374_G_C | 12 | 228374 | 1.83e-07 | Yes |
| 3545271_chrVI_76442_G_A | 6 | 76442 | 4.56e-15 | Yes |
| 7807333_chrXII_562305_C_G | 12 | 562305 | 1.11e-170 | Yes |
| 11055361_chrXV_1023392_T_G | 15 | 1023392 | 2.33e-07 | Yes |
| 1757949_chrIV_397927_G_A | 4 | 397927 | 1.64e-51 | Yes |
| 11517118_chrXVI_393858_A_G | 16 | 393858 | 1.75e-128 | Yes |
| 5558748_chrIX_166175_C_T | 9 | 166175 | 3.30e-42 | Yes |
| 1171626_chrIII_128224_T_C | 3 | 128224 | 1.42e-48 | Yes |
| 11578735_chrXVI_455475_T_A | 16 | 455475 | 7.08e-94 | Yes |
| 1202358_chrIII_158956_T_C | 3 | 158956 | 1.51e-47 | Yes |
| 2074955_chrIV_714933_T_C | 4 | 714933 | 9.69e-25 | Yes |
| 7330660_chrXII_85632_T_G | 12 | 85632 | 5.07e-13 | Yes |
| 4023350_chrVII_284360_A_G | 7 | 284360 | 2.19e-101 | Yes |
| 9185900_chrXIII_862695_C_T | 13 | 862695 | 6.24e-57 | Yes |
| 1107706_chrIII_64304_C_T | 3 | 64304 | 4.07e-16 | Yes |
| 1581197_chrIV_221175_G_A | 4 | 221175 | 7.77e-32 | Yes |
| 6611885_chrXI_33673_G_A | 11 | 33673 | 7.92e-23 | Yes |
| 8623529_chrXIII_300324_G_C | 13 | 300324 | 0.00e+00 | Yes |
| 10414113_chrXV_382144_C_T | 15 | 382144 | 1.53e-205 | Yes |
| 5928463_chrX_96002_G_T | 10 | 96002 | 6.04e-193 | Yes |
| 3567664_chrVI_98835_T_A | 6 | 98835 | 3.90e-21 | Yes |
| 2440545_chrIV_1080523_C_T | 4 | 1080523 | 2.28e-36 | Yes |
| 7996016_chrXII_750988_C_T | 12 | 750988 | 2.41e-253 | Yes |
| 6848038_chrXI_269826_A_G | 11 | 269826 | 1.64e-24 | Yes |
| 760082_chrII_529864_G_A | 2 | 529864 | 1.22e-114 | Yes |
| 925586_chrII_695368_T_C | 2 | 695368 | 6.30e-24 | Yes |
| 6594950_chrXI_16738_T_C | 11 | 16738 | 6.50e-15 | Yes |
| 5678919_chrIX_286346_G_A | 9 | 286346 | 1.79e-66 | Yes |
| 9435376_chrXIV_187740_T_C | 14 | 187740 | 3.24e-68 | Yes |
| 11739349_chrXVI_616089_G_A | 16 | 616089 | 6.69e-20 | Yes |
| 3161012_chrV_269057_A_G | 5 | 269057 | 4.73e-60 | Yes |
| 9167252_chrXIII_844047_C_T | 13 | 844047 | 1.99e-49 | Yes |
| 10551485_chrXV_519516_C_G | 15 | 519516 | 3.83e-75 | Yes |
| 5093312_chrVIII_263382_T_C | 8 | 263382 | 1.59e-31 | Yes |
| 138823_chrI_138823_A_G | 1 | 138823 | 1.04e-44 | Yes |
| 7921913_chrXII_676885_G_A | 12 | 676885 | 0.00e+00 | Yes |
| 5303814_chrVIII_473884_A_G | 8 | 473884 | 1.45e-26 | Yes |
| 3885183_chrVII_146193_A_G | 7 | 146193 | 0.00e+00 | Yes |
| 9462469_chrXIV_214833_G_A | 14 | 214833 | 8.34e-103 | Yes |
| 2413305_chrIV_1053283_G_A | 4 | 1053283 | 3.12e-42 | Yes |
| 935324_chrII_705106_A_T | 2 | 705106 | 5.91e-25 | Yes |
| 11020152_chrXV_988183_G_A | 15 | 988183 | 8.40e-13 | Yes |
| 6324127_chrX_491666_G_A | 10 | 491666 | 5.22e-81 | Yes |
| 2248498_chrIV_888476_T_C | 4 | 888476 | 4.55e-43 | Yes |
| 6997404_chrXI_419192_A_G | 11 | 419192 | 3.12e-41 | Yes |
| 6751527_chrXI_173315_G_C | 11 | 173315 | 1.69e-42 | Yes |
| 9536458_chrXIV_288822_C_T | 14 | 288822 | 3.39e-202 | Yes |
| 8151459_chrXII_906431_T_C | 12 | 906431 | 4.57e-25 | Yes |
| 8493990_chrXIII_170785_T_A | 13 | 170785 | 7.79e-260 | Yes |
| 1342500_chrIII_299098_C_T | 3 | 299098 | 2.00e-07 | Yes |
| 8847982_chrXIII_524777_C_A | 13 | 524777 | 3.13e-25 | Yes |
| 5979363_chrX_146902_A_G | 10 | 146902 | 1.49e-155 | Yes |
| 10988277_chrXV_956308_G_T | 15 | 956308 | 7.30e-15 | Yes |
| 11217478_chrXVI_94218_T_C | 16 | 94218 | 2.74e-20 | Yes |
| 6379661_chrX_547200_C_T | 10 | 547200 | 3.97e-105 | Yes |
| 3520086_chrVI_51257_A_G | 6 | 51257 | 8.69e-10 | Yes |
| 10864545_chrXV_832576_G_A | 15 | 832576 | 1.18e-28 | Yes |
| 10244782_chrXV_212813_T_C | 15 | 212813 | 2.15e-316 | Yes |
| 6646084_chrXI_67872_C_G | 11 | 67872 | 6.25e-70 | Yes |
| 7011842_chrXI_433630_T_C | 11 | 433630 | 3.12e-31 | Yes |
| 5515513_chrIX_122940_A_T | 9 | 122940 | 1.84e-55 | Yes |
| 9901193_chrXIV_653557_C_T | 14 | 653557 | 3.50e-89 | Yes |
| 8836630_chrXIII_513425_A_T | 13 | 513425 | 1.38e-16 | Yes |
| 4726124_chrVII_987134_T_A | 7 | 987134 | 2.34e-22 | Yes |
| 188190_chrl_188190_T_C | 1 | 188190 | 3.19e-130 | Yes |
| 1598848_chrIV_238826_G_T | 4 | 238826 | 3.91e-32 | Yes |
| 4581196_chrVII_842206_T_C | 7 | 842206 | 8.72e-82 | Yes |
| 6256867_chrX_424406_A_T | 10 | 424406 | 2.78e-41 | Yes |
| 1861689_chrIV_501667_A_C | 4 | 501667 | 5.74e-29 | Yes |
| 5968656_chrX_136195_G_A | 10 | 136195 | 8.97e-212 | Yes |
| 6280789_chrX_448328_C_T | 10 | 448328 | 2.54e-41 | Yes |
| 602948_chrII_372730_G_A | 2 | 372730 | 1.32e-88 | Yes |
| 2419116_chrIV_1059094_C_T | 4 | 1059094 | 4.86e-42 | Yes |
| 6452525_chrX_620064_T_C | 10 | 620064 | 2.52e-84 | Yes |
| 1414245_chrIV_54223_C_T | 4 | 54223 | 4.54e-27 | Yes |
| 9300295_chrXIV_52659_G_T | 14 | 52659 | 2.60e-03 | No |
| 9773211_chrXIV_525575_T_C | 14 | 525575 | 0.00e+00 | Yes |
| 6945298_chrXI_367086_A_T | 11 | 367086 | 8.70e-51 | Yes |
| 6136044_chrX_303583_C_T | 10 | 303583 | 1.96e-08 | Yes |
| 10262035_chrXV_230066_C_T | 15 | 230066 | 4.31e-263 | Yes |
| 8293957_chrXII_1048929_G_T | 12 | 1048929 | 5.47e-02 | No |
| 4070183_chrVII_331193_A_G | 7 | 331193 | 1.46e-59 | Yes |
| 3847228_chrVII_108238_G_C | 7 | 108238 | 2.46e-262 | Yes |
| 1728757_chrIV_368735_G_A | 4 | 368735 | 3.65e-49 | Yes |
| 1989747_chrIV_629725_G_A | 4 | 629725 | 2.30e-26 | Yes |
| 2221279_chrIV_861257_A_G | 4 | 861257 | 2.41e-33 | Yes |
| 1606765_chrIV_246743_G_C | 4 | 246743 | 2.81e-35 | Yes |
| 3252364_chrV_360409_C_A | 5 | 360409 | 2.41e-47 | Yes |
| 11014104_chrXV_982135_A_T | 15 | 982135 | 1.46e-13 | Yes |
| 1467219_chrIV_107197_G_T | 4 | 107197 | 5.23e-13 | Yes |
| 9795044_chrXIV_547408_G_A | 14 | 547408 | 3.81e-222 | Yes |
| 8084909_chrXII_839881_G_A | 12 | 839881 | 7.77e-50 | Yes |
| 5214936_chrVIII_385006_G_A | 8 | 385006 | 2.08e-24 | Yes |
| 3965142_chrVII_226152_C_T | 7 | 226152 | 0.00e+00 | Yes |
| 11174690_chrXVI_51430_C_T | 16 | 51430 | 8.16e-18 | Yes |
| 8212296_chrXII_967268_A_G | 12 | 967268 | 2.20e-12 | Yes |
| 6297026_chrX_464565_C_A | 10 | 464565 | 6.08e-47 | Yes |
| 4878229_chrVIII_48299_T_C | 8 | 48299 | 2.41e-97 | Yes |
| 5455388_chrIX_62815_A_G | 9 | 62815 | 3.39e-35 | Yes |
| 3037458_chrV_145503_A_G | 5 | 145503 | 4.44e-33 | Yes |
| 2693492_chrIV_1333470_C_T | 4 | 1333470 | 3.48e-17 | Yes |
| 3968793_chrVII_229803_G_A | 7 | 229803 | 4.98e-318 | Yes |
| 8933051_chrXIII_609846_C_T | 13 | 609846 | 7.40e-69 | Yes |
| 707172_chrII_476954_C_T | 2 | 476954 | 2.94e-135 | Yes |
| 173222_chrI_173222_A_C | 1 | 173222 | 8.53e-73 | Yes |
| 7820825_chrXII_575797_G_T | 12 | 575797 | 9.97e-252 | Yes |
| 6402295_chrX_569834_C_T | 10 | 569834 | 3.88e-84 | Yes |
| 939170_chrII_708952_T_C | 2 | 708952 | 7.15e-24 | Yes |
| 6085025_chrX_252564_T_C | 10 | 252564 | 8.19e-31 | Yes |
| 4794687_chrVII_1055697_A_G | 7 | 1055697 | 9.89e-05 | Yes |
| 7205226_chrXI_627014_C_T | 11 | 627014 | 1.41e-59 | Yes |
| 11695724_chrXVI_572464_T_A | 16 | 572464 | 4.51e-47 | Yes |
| 3716953_chrVI_248124_C_T | 6 | 248124 | 7.87e-06 | Yes |
| 2819033_chrIV_1459011_C_T | 4 | 1459011 | 2.19e-07 | Yes |
| 1941203_chrIV_581181_T_C | 4 | 581181 | 8.43e-28 | Yes |
| 337341_chrII_107123_A_T | 2 | 107123 | 1.62e-04 | No |
| 4572794_chrVII_833804_C_A | 7 | 833804 | 2.43e-71 | Yes |
| 8040064_chrXII_795036_C_A | 12 | 795036 | 5.11e-112 | Yes |
| 5689026_chrIX_296453_T_A | 9 | 296453 | 4.06e-53 | Yes |
| 450642_chrII_220424_A_C | 2 | 220424 | 3.39e-34 | Yes |
| 9008894_chrXIII_685689_T_C | 13 | 685689 | 8.82e-119 | Yes |
| 8661636_chrXIII_338431_A_G | 13 | 338431 | 7.00e-253 | Yes |
| 60384_chrI_60384_C_T | 1 | 60384 | 1.01e-122 | Yes |
| 8519173_chrXIII_195968_G_A | 13 | 195968 | 4.96e-289 | Yes |
| 3227559_chrV_335604_T_C | 5 | 335604 | 1.10e-40 | Yes |
| 6301141_chrX_468680_G_A | 10 | 468680 | 1.07e-55 | Yes |
| 11027149_chrXV_995180_A_G | 15 | 995180 | 5.09e-09 | Yes |
| 5050085_chrVIII_220155_T_C | 8 | 220155 | 1.27e-99 | Yes |
| 6036897_chrX_204436_C_A | 10 | 204436 | 1.10e-48 | Yes |
| 1393233_chrIV_33211_A_G | 4 | 33211 | 1.86e-25 | Yes |
| 1694415_chrIV_334393_T_C | 4 | 334393 | 4.63e-41 | Yes |
| 4913863_chrVIII_83933_G_A | 8 | 83933 | 5.06e-313 | Yes |
| 3633940_chrVI_165111_A_G | 6 | 165111 | 1.07e-16 | Yes |
| 1441607_chrIV_81585_T_C | 4 | 81585 | 2.08e-14 | Yes |
| 130776_chrI_130776_C_G | 1 | 130776 | 3.76e-37 | Yes |
| 11204417_chrXVI_81157_G_A | 16 | 81157 | 4.19e-19 | Yes |
| 281133_chrII_50915_C_T | 2 | 50915 | 1.25e-05 | Yes |
| 8951620_chrXIII_628415_G_A | 13 | 628415 | 1.25e-87 | Yes |
| 7528822_chrXII_283794_C_T | 12 | 283794 | 2.80e-35 | Yes |
| 2767557_chrIV_1407535_A_G | 4 | 1407535 | 4.30e-22 | Yes |

<sup>1</sup>G x E: whether the p value reaches multiple testing threshold

## References

1. Forsberg, S. K. G., Bloom, J. S., Sadhu, M. J., Kruglyak, L. & Carlborg, Ö. Accounting for genetic interactions improves modeling of individual quantitative trait phenotypes in yeast. Nat. Genet. 49, 497–503 (2017).

2. Juenger, T. E., Sen, S., Stowe, K. A. & Simms, E. L. Epistasis and genotype-environment interaction for quantitative trait loci affecting flowering time in Arabidopsis thaliana. Genetica 123, 87–105 (2005).

3. Carlborg, O., Jacobsson, L., Ahgren, P., Siegel, P. & Andersson, L. Epistasis and the release of genetic variation during long-term selection. Nat. Genet. 38, 418–420 (2006).

4. Sasaki, E., Zhang, P., Atwell, S., Meng, D. & Nordborg, M. Missing G x E Variation Controls Flowering Time in Arabidopsis thaliana. PLOS Genet. 11, e1005597 (2015).

5. Smith, A. N., Miller, L.-A., Radice, G., Ashery-Padan, R. & Lang, R. A. Stage-dependent modes of Pax6-Sox2 epistasis regulate lens development and eye morphogenesis. Development 136, 3377–3377 (2009).

6. Kerwin, R. E. et al. Epistasis × environment interactions among Arabidopsis thaliana glucosinolate genes impact complex traits and fitness in the field. New Phytol. 215, 1249–1263 (2017).

7. Bhatia, A. et al. Yeast Growth Plasticity Is Regulated by Environment-Specific Multi-QTL Interactions. G3 4, 769–777 (2014).

8. Yadav, A. & Sinha, H. Gene-gene and gene-environment interactions in complex traits in yeast. Yeast 35, 403–416 (2018).

9. Yadav, A., Dhole, K. & Sinha, H. Differential regulation of cryptic genetic variation shapes the genetic interactome underlying complex traits. Genome Biol. Evol. 8, 3559–3573 (2016). doi:10.1093/gbe/evw258

10. Flynn, K. M., Cooper, T. F., Moore, F. B. G. & Cooper, V. S. The Environment Affects Epistatic Interactions to Alter the Topology of an Empirical Fitness Landscape. PLoS Genet. 9, e1003426 (2013).

11. Li, C. & Zhang, J. Multi-environment fitness landscapes of a tRNA gene. *Nat*. Ecol. Evol. 2, 1025–1032 (2018).

12. de Vos, M. G. J., Poelwijk, F. J., Battich, N., Ndika, J. D. T. & Tans, S. J. Environmental Dependence of Genetic Constraint. PLoS Genet. 9, e1003580 (2013).

13. Remold, S. K. & Lenski, R. E. Pervasive joint influence of epistasis and plasticity on mutational effects in Escherichia coli. Nat. Genet. 36, 423–426 (2004).

14. Lee, J. T., Coradini, A. L. V., Shen, A. & Ehrenreich, I. M. Layers of Cryptic Genetic Variation Underlie a Yeast Complex Trait. Genetics 211, 1469–1482 (2019).

15. Hou, J., van Leeuwen, J., Andrews, B. J. & Boone, C. Genetic Network Complexity Shapes Background-Dependent Phenotypic Expression. Trends Genet. 34, 578–586 (2018).

16. Mullis, M. N., Matsui, T., Schell, R., Foree, R. & Ehrenreich, I. M. The complex underpinnings of genetic background effects. Nat. Commun. 9, 3548 (2018).

17. Lee, J. T., Taylor, M. B., Shen, A. & Ehrenreich, I. M. Multi-locus Genotypes Underlying Temperature Sensitivity in a Mutationally Induced Trait. PLOS Genet. 12, e1005929 (2016).

18. Bandyopadhyay, S. et al. Rewiring of Genetic Networks in Response to DNA Damage. Science 330, 1385–1389 (2010).

19. Kuzmin, E. et al. Systematic analysis of complex genetic interactions. Science 360, eaao1729 (2018).

20. Leung, G. P., Aristizabal, M. J., Krogan, N. J. & Kobor, M. S. Conditional Genetic Interactions of RTT107, SLX4, and HRQ1 Reveal Dynamic Networks upon DNA Damage in S. cerevisiae. G3 4, 1059–1069 (2014).

21. Bloom, J. S. et al. Genetic interactions contribute less than additive effects to quantitative trait variation in yeast. Nat. Commun. 6, 8712 (2015).

22. Albert, F. W., Bloom, J. S., Siegel, J., Day, L. & Kruglyak, L. Genetics of trans-regulatory variation in gene expression. Elife 7, e35471 (2018).

23. Bloom, J. S., Ehrenreich, I. M., Loo, W. T., Lite, T.-L. V. & Kruglyak, L. Finding the sources of missing heritability in a yeast cross. Nature 494, 234– 237 (2013).

24. Gibson, G. Decanalization and the origin of complex disease. Nat. Rev. Genet. 10, 134–140 (2009).

25. Paaby, A. B. & Rockman, M. V. Cryptic genetic variation: evolution’s hidden substrate. Nat. Rev. Genet. 15, 247–258 (2014).

26. Hayden, E. J., Ferrada, E. & Wagner, A. Cryptic genetic variation promotes rapid evolutionary adaptation in an RNA enzyme. Nature 474, 92–95 (2011).

27. Queitsch, C., Sangster, T. A. & Lindquist, S. Hsp90 as a capacitor of phenotypic variation. Nature 417, 618–624 (2002).

28. Rutherford, S. L. & Lindquist, S. Hsp90 as a capacitor for morphological evolution. Nature 396, 336–342 (1998).

29. Dworkin, I., Palsson, A., Birdsall, K. & Gibson, G. Evidence that Egfr contributes to cryptic genetic variation for photoreceptor determination in natural populations of Drosophila melanogaster. Curr. Biol. 13, 1888–93 (2003).

30. Miyajima, I. et al. GPA1, a haploid-specific essential gene, encodes a yeast homolog of mammalian G protein which may be involved in mating factor signal transduction. Cell 50, 1011–1019 (1987).

31. Bourgarel, D., Nguyen, C.-C. & Bolotin-Fukuhara, M. HAP4, the glucose-repressed regulated subunit of the HAP transcriptional complex involved in the fermentation-respiration shift, has a functional homologue in the respiratory yeast Kluyveromyces lactis. Mol. Microbiol. 31, 1205–1215 (1999).

32. Zhang, L. & Hach, A. Molecular mechanism of heme signaling in yeast: the transcriptional activator Hap1 serves as the key mediator. Cell. Mol. Life Sci. 56, 415–426 (1999).

33. Creusot, F., Verdière, J., Gaisne, M. & Slonimski, P. P. CYP1 (HAP1) regulator of oxygen-dependent gene expression in yeast. J. Mol. Biol. 204, 263–276 (1988).

34. Sharma, S. et al. Yeast Kre33 and human NAT10 are conserved 18S rRNA cytosine acetyltransferases that modify tRNAs assisted by the adaptor Tan1/THUMPD1. Nucleic Acids Res. 43, 2242–58 (2015).

35. Wickner, R. B. MKT1, a nonessential Saccharomyces cerevisiae gene with a temperature-dependent effect on replication of M2 double-stranded RNA. J. Bacteriol. 169, 4941–4945 (1987).

36. Tanaka, K., Lin, B. K., Wood, D. R. & Tamanoi, F. IRA2, an upstream negative regulator of RAS in yeast, is a RAS GTPase-activating protein. Proc. Natl. Acad. Sci. 88, 468–472 (1991).

37. Smith, E. N. & Kruglyak, L. Gene–Environment Interaction in Yeast Gene Expression. PLoS Biol. 6, e83 (2008).

38. Lewis, J. A., Broman, A. T., Will, J. & Gasch, A. P. Genetic architecture of ethanol-responsive transcriptome variation in Saccharomyces cerevisiae strains. Genetics 198, 369–82 (2014).

39. Boyle, E. A., Li, Y. I. & Pritchard, J. K. An Expanded View of Complex Traits: From Polygenic to Omnigenic. Cell 169, 1177–1186 (2017).

40. Zan, Y. et al. Artificial Selection Response due to Polygenic Adaptation from a Multilocus, Multiallelic Genetic Architecture. Mol. Biol. Evol. 2, 7–10 (2017).

41. Sheng, Z., Pettersson, M. E., Honaker, C. F., Siegel, P. B. & Carlborg, Ö. Standing genetic variation as a major contributor to adaptation in the Virginia chicken lines selection experiment. Genome Biol. 16, 219 (2015).

42. Zeileis, A. & Hothorn, T. Diagnostic checking in regression relationships. R News 2, 7–10 (2002).

43. R Core Team. R Core Team, 2015 R: A Language and Environment for Statistical Computing. R Found. Stat. Comput. Vienna Austria URL: https://www.R-project.org/. R Foundation for Statistical Computing Vienna Austria (2015). doi:ISBN 3-900051-07-0

44. Yu, G., Chen, Y. & Guo, Y. Design of integrated system for heterogeneous network query terminal. *J*. Comput. Appl. 29, 2191–2193 (2009).

